# Efficient Simulations of Membrane and Solvent Asymmetry With Flat-Bottom Restraints

**DOI:** 10.1101/2023.05.26.542460

**Authors:** Denys Biriukov, Matti Javanainen

## Abstract

The routinely employed periodic boundary conditions complicate molecular simulations of physiologically relevant asymmetric lipid membranes together with their distinct solvent environments. Therefore, separating the extracellular fluid from its cytosolic counterpart has often been performed using a costly double-bilayer setup. Here, we demonstrate that the lipid membrane and solvent asymmetry can be efficiently modeled with a single lipid bilayer by applying a flat-bottom potential to ions and other solute molecules, thereby restraining them to only interact with its relevant leaflet. We carefully optimized the parameters of the suggested method so that the results obtained using the flat-bottom and double-bilayer approaches become mutually indistinguishable. Then, we apply the flat-bottom approach to lipid bilayers with various compositions and solvent environments, covering ions and cationic peptides to validate the approach in a realistic use case. We also discuss the possible limitations of the method as well as its computational efficiency and provide a step-by-step guide on how to set up such simulations in a straightforward manner.

## Introduction

Biological membranes, such as the plasma membrane and membranes encapsulating the cellular organelles, have long been known to be asymmetric in their compositions.^1^ Lipidomics studies have resolved the asymmetry of the membrane leaflets in terms of lipid head groups^2^ as well as chain length and lipid saturation.^3^ This asymmetry leads to distinct biophysical properties of the membrane leaflets that are crucial for many cellular functions. ^4^ Thus, the lipid asymmetry is maintained and tightly regulated by protein machinery.^5^

Not only the lipid composition but also the solvent environments of the two leaflets of biomembranes can differ significantly.^6^ For example, in the case of the plasma membrane, the extracellular fluid and the cytosol contain *≈*140 mM of Na^+^ and K^+^, respectively. These charges are mainly neutralized by Cl*^−^* and some hydrogencarbonate in the extracellular fluid and by negatively charged organic phosphates and acids as well as proteins in the cytosol. Divalent cations are present in smaller amounts, Ca^2+^ mainly in the extracellular fluid and Mg^2+^ primarily in the cytoplasmic matrix. As the membranes are generally impermeable to ions, numerous cellular processes can be driven by the membrane potential resulting from the ion imbalances.^6^

Molecular dynamics (MD) simulations of biomolecules and their assemblies have become an essential tool in the fields of physical chemistry, biophysics, biochemistry, and structural biology. This development is supported by steadily increasing computational power, which allows the sampling of even more realistic systems in reasonable time scales.^7, 8^ However, despite the compositional complexity of biomembranes being regularly incorporated in the simulation models, the leaflet asymmetry is less often considered.^3, 9–11^ One of the reasons for this omission is that whereas bilayers with symmetric lipid composition are readily simulated in their native tensionless state, this is tedious to achieve for asymmetric leaflets. The most common approach is to match the areas per lipid of the two leaflets,^12^ yet this has been shown to be potentially inaccurate.^13^ Instead, the tension of each leaflet should be eliminated by the membrane setup,^13^ which is at least aided if not eliminated by regular flip–flops of membrane components such as cholesterol.^14, 15^

Another requirement for the simulations of realistic membranes is that the solvent environments to which the asymmetric leaflets are exposed should differ. However, this is also a challenge for MD simulations that commonly exploit periodic boundary conditions (PBCs) to eliminate boundary effects and ensure the conservation of linear momentum. Due to PBC, the solvent in contact with the leaflets in a simple single-bilayer setup is continuous across the simulation box. Thus, in order to create two different solvent environments interacting with two corresponding leaflets, a double-bilayer setup is required (see examples on the top row of Fig. 1).^16^ In this setup, the two bilayers confine two distinct solvents, and the bilayer leaflets are exposed to different solvent environments.

**Figure 1:**
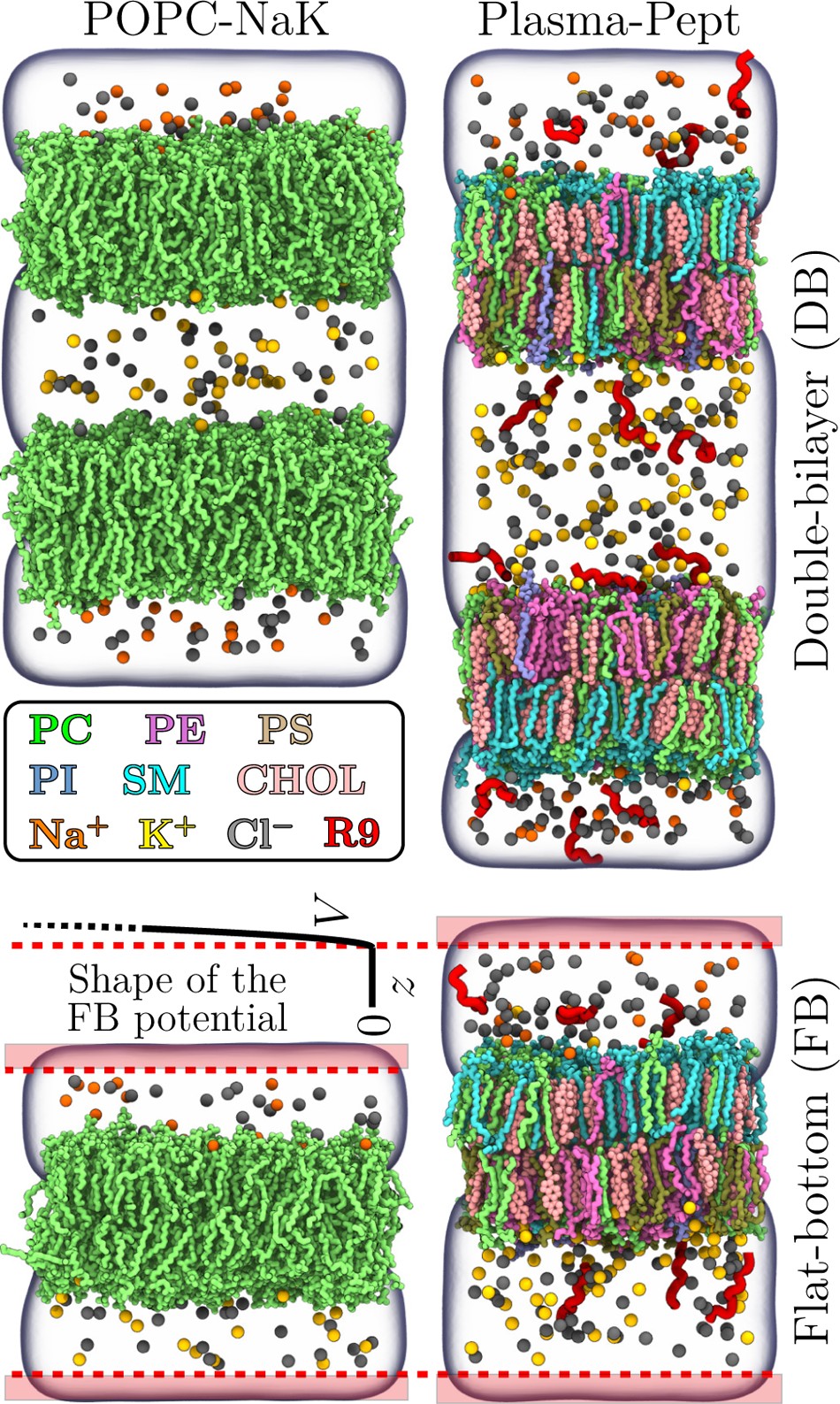
Snapshots of two example systems with the flat-bottom and double-bilayer setups. The coloring is provided in the legend. Water is rendered as a transparent volume. Lipid hydrogens are not rendered for clarity. The flat-bottom is schematically visualized by the shape and position of the potential *V*, drawn for the “Plasma-Pept-FB” system. The red shaded areas demonstrate the zones of extra water where ions or peptides cannot enter. The flat-bottom potential becomes effective at the dashed red line, *i.e.*, 0.3 nm from the box edge.

The double-bilayer setup has been used to generate electrostatic potential with explicition imbalance,^16, 17^ study the effect of natural Na^+^/K^+^ imbalance on membrane properties,^18^ and investigate the behavior of a charged peptide.^19^ However, the double-bilayer setup has its own drawbacks as well. The doubling of the system size also increases the computational cost in theory by a factor of two, and even more when long-range non-bonded interactions are involved.^20, 21^ While this can be a minor setback for lipid-only membranes, it becomes unbearable for simulations of large membrane–protein complexes. Moreover, any interactions of, *e.g.*, proteins or other molecules with the bilayer should occur simultaneously with both bilayers of the double-bilayer setup. Otherwise, the perturbations caused by this interaction can be partially eliminated by the non-interacting bilayer. As an example, the condensation of one bilayer by, *e.g.*, bound ions would be opposed by the other bilayer with typically very small compressibility.

On some occasions, the limitations with modeling the lipid asymmetry can be surmounted by including the membrane potential *via* an applied external electric field instead of explicitly modeled ion imbalance. While these two approaches have similar effects on the bilayer properties^22^ as well as on the energetics of membrane pore formation,^23^ this is not always sufficient. If the role of ions or other charged molecules is not only to induce the potential, such as in the case of specific protein–ion interactions,^24, 25^ the electric field approach is clearly inadequate. Thereby, there is a demand for an efficient approach for modeling separate solvent environments in contact with the membrane leaflets without adding significant computational overhead.

Flat-bottom potentials—or flat-bottom restraints—are implemented in multiple modern MD engines. They have been used, for instance, to extract osmotic coefficients,^26^ to include NMR restraints into MD refinement of proteins and nucleic acids,^27^ to adjust local concentrations of ions,^28^ and to induce pores that facilitate lipid flip–flops across bilayer leaflets.^29^ Here, we demonstrate that flat-bottom potentials can be used to maintain the ionic asymmetry across a single lipid bilayer in molecular dynamics simulations. The idea of the approach is to keep ions or other molecules in the vicinity of corresponding leaflets. The flat-bottom approach can be potentially expanded to other soluble molecules of physiological or engineering importance, such as drugs and other small molecules as well as cytosolic proteins. Its potential use cases that we identified are to either model physiological ion imbalances across bilayers or to facilitate the sampling of adsorption/desorption events of, *e.g.*, peptides onto the bilayer. The latter can be achieved by having different adsorbents on the two sides of the bilayer, which effectively halves the required simulation time when multiple adsorbents are sampled. Alternatively, if the same adsorbent is simulated on both sides of the membrane at a well-defined concentration, the sampling gained in a single simulation is effectively doubled. Moreover, the approach is independent of the used force fields and simulation software.

The open question we tackle in this work is whether the behavior of ions, charged peptides, and membrane lipids is affected by the flat-bottom approach or whether it is indistinguishable from a significantly more costly double-bilayer setup. There are indeed potential pitfalls to consider. Large imbalances in ionic strength (or, essentially, activity or osmotic coefficients) across the two sides of the membrane may lead to a non-zero osmotic gradient, which can displace the membrane with respect to the flat-bottom potential and thus alter the concentrations of the salts on the two sides of the membrane. However, the same limitations naturally apply to double-bilayer setups, with the only difference being that this is not often realized within the timescale of typical MD simulations. Here, we carefully evaluate the impact of such effects on the cases of ion and peptide imbalance together with symmetric and asymmetric membranes.

## Methods

### Flat-bottom potential

In the GROMACS simulation engine^30, 31^—used throughout this work—a flat-bottom potential acting on a layer with a constant *z* coordinate is implemented with a quadratic form

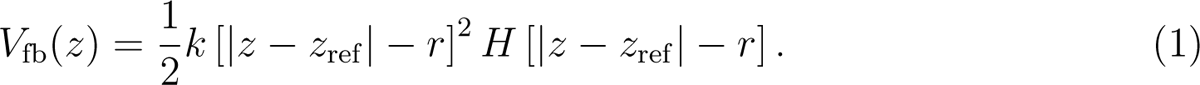

Here, *z* is the coordinate of the restrained atom, whereas the reference coordinate, *z*_ref_, is at the center of the flat-bottom potential. The width of the flat-bottom potential is *r*, and in the case of the regular flat-bottom potential, the potential is zero across a length of 2 *× r* centered at *z*_ref_. However, here we use the inverted form of the potential so that it is zero except for a region 2 *× r* wide centered at *z*_ref_. The force constant *k* controls how steeply the potential grows. The flat-bottom nature is provided by the Heaviside function *H*, which is zero elsewhere but one when the distance of *z* from the reference *z*_ref_ is smaller than *r*.

### Water

We used simple aqueous salt solutions to first calibrate the two parameters of the flat-bottom potential, *r* and *k*, which will be then used for the lipid bilayer simulations (see below). To this end, we set up simulation boxes with *∼*2000 water molecules and 108 pairs of Na^+^ and Cl*^−^*. The boxes were elongated with dimensions of 3*×*3*×*8 nm^3^. We evaluated which values of *r* and *k* would ensure that no ions would cross the box edge where the flatbottom potential was centered (*z*_ref_ = 0). Notably, the smaller the *r* value, the smaller the region perturbed by the flat-bottom potential is. We performed simulations with the force constant *k* set to 100, 1000, 10,000, or 100,000 kJ*·*mol*^−^*^1^*·*nm*^−^*^2^ and with *r* set to 0.1, 0.2, 0.3, 0.4, or 0.5 nm. The long dimension of the simulation box was extended by the value of *r* so that there was always an 8 nm-wide unperturbed region. We simulated the systems for 100 ns and calculated the numbers of ion crossings across the flat-bottom potential.

Despite using as small as possible value for *r*, a small fraction of the simulation box is still perturbed; ions (or other molecules to which flat-bottom potential is applied) are depleted in this region, slightly increasing their concentration in the unperturbed region. This raises a question on how much *extra water* should be added to the system to have the size of its unperturbed region match that of a system without a flat-bottom potential applied, *i.e.*, to describe the target concentration of ions (or other molecules) in the unperturbed region. Notably, this is often not necessary for cases involving larger molecules, where only one such molecule interacts with a bilayer leaflet, and the correction to experimental concentration can be done analytically after the simulation. However, in the case of ions, it is crucial to match the concentrations between the simulation and the reference experiment.^32^ To study how much extra water should be included in the presence of flat-bottom restraints, we simulated the water box with varying values of the long dimension (8.1–8.8 nm with a step of 0.1 nm) with the *r* and *k* values optimized using the simulations described above. We calculated the Na^+^ density profiles, averaged the density in the unperturbed region, and compared these values to the one extracted from a simulation without a flat-bottom potential and with a long dimension of 8 nm.

### Lipid bilayers

To evaluate the viability of the flat-bottom approach, we set up multiple test cases involving different lipid compositions and different salts, including also more complex molecules in the form of cationic nona-arginine peptides, present on one or two sides of a membrane. As the simplest example, we first considered a POPC bilayer consisting of 308 lipids divided equally between the two leaflets. In addition, a more complex bilayer whose composition mimicked the asymmetry of the plasma membrane^3^ was used, see Table S1 in the Supporting Information (SI) for the detailed composition. These membranes were set up using CHARMM-GUI^33, 34^ with outputs in GROMACS formats.^35^ The membranes were excessively hydrated by 50 water molecules per lipid, and different salts were placed on the two sides of the membrane. For each system, a corresponding double-bilayer setup was also set up by rotating and translating the original bilayer system using the tool gmx editconf of GROMACS, followed by concatenating these two systems into one. Finally, an extra layer of water was included in the flat-bottom systems based on the findings of the simulations with aqueous salt solutions described above. Thus, for the two membrane systems, the number of lipids and ions was doubled, whereas with water, the ratio is slightly smaller than two due to the extra solvent (see “Water” above). Snapshots of two selected systems using both the flat-bottom and double-bilayer approaches are shown in Fig. 1. All simulated lipid bilayer systems are listed in Table 1. The input and output data from all simulations are openly available in Zenodo at DOI:.

**Table 1:**
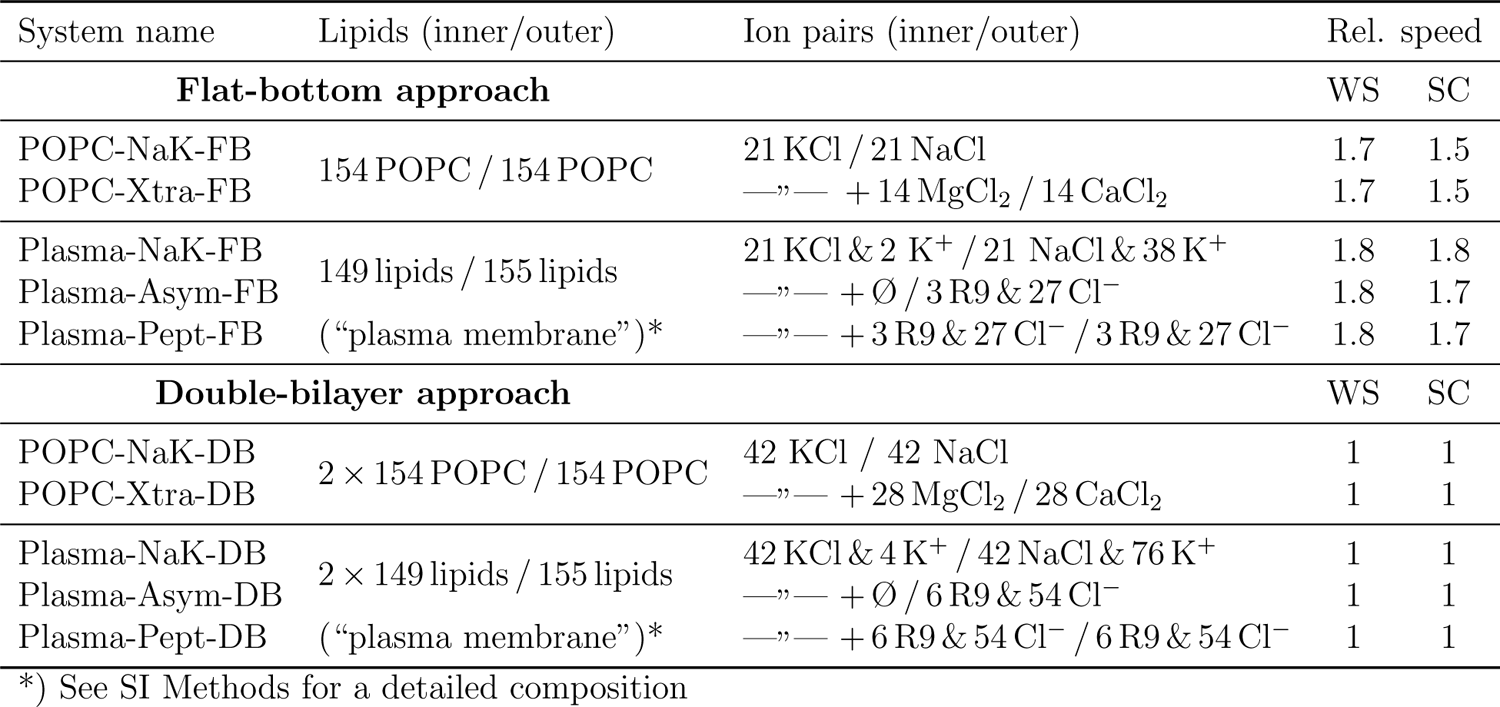
Summary of simulated systems with their lipid and solution compositions with respect to inner (cytosolic) and outer (extracellular) leaflet/solution compartments. Relative speed is given as a ratio of the ns/day values obtained for the flat-bottom and double-bilayer approaches using a typical workstation setup with a powerful GPU (“WS”) and a CPU-only supercomputer (“SC”). Details of their configurations are provided in the Supporting Information. The standard deviation in relative speed is less than one percent and thus not reported.

The CHARMM36 force field^36, 37^ was used for lipids and the CHARMM-specific TIP3P model for water.^38, 39^ For the nona-arginine (R9) peptides, we used the scaled-charge pros-ECCo75 force field based on CHARMM36m.^40, 41^ The latter was necessary to properly sample binding/unbinding events of R9 peptides and ensure sufficient convergence of flat-bottom and double-bilayer simulations.^32, 41^ All membranes were first energy-minimized and equilibrated using the standard protocol obtained from CHARMM-GUI,^35^ after which were simulated for 1 *µ*s using a leap-frog integrator with a time step of 2 fs with the GROMACS simulation engine,^30^ versions 2021, 2021.5, and 2022.3. Buffered Verlet lists^42^ were used to keep track of atomic neighbors. Long-range electrostatics were handled using smooth particle mesh Ewald (PME) algorithm.^20, 43^ Lennard-Jones potential was cut-off at 1.2 nm with the forces switched to zero starting at a distance of 1.0 nm. The Nośe–Hoover thermostat ^44, 45^ with a target temperature of 310 K and a coupling time constant of 1 ps was applied separately to the lipids and the solvent. The Parrinello–Rahman barostat^46^ with a reference pressure of 1 bar, coupling time constant of 5 ps, and compressibility of 4.5*×*10*^−^*^5^ bar*^−^*^1^ was applied semi-isotropically to the membrane plane (*x* and *y* dimensions) and normal to it (*z*), respecting the symmetry of the system. Bonds in the lipid molecules involving hydrogen atoms were constrained using P-LINCS,^47, 48^ whereas the geometry of water was constrained using SETTLE.^49^ The trajectories were written every 100 ps, and the first 100 ns were omitted from all analyses. In the flat-bottom simulations, we applied the restraints to all ions and all heavy atoms of the peptides. For selected systems, we also checked applying the restraints only to cations (*i.e.* we excluded the restraints from Cl*^−^* anions), which served as an additional test of the flat-bottom approach. We used the parameters optimized with the salt solution simulations, namely *r*=0.3 nm, *k*=10,000 kJ*·*mol*^−^*^1^*·*nm*^−^*^2^, and an extra layer of water with a thickness of *∼*0.3 nm on each side of the membrane.

The area per lipid (APL) was extracted by dividing the simulation box area by the number of non-cholesterol lipids in one leaflet. Membrane thickness (*D*_P_*_−_*_P_) was evaluated as the inter-leaflet distance of the average position of phosphorus atoms. The P–N vector tilt angle describes the angle between the membrane normal and the vector connecting the phosphorus and nitrogen atoms in the choline head group of phosphatidylcholine (PC) lipids. The number density profiles, water dipole orientation, and deuterium order parameters were calculated using gmx density, gmx h2order, and gmx order tools, respectively. In the case of double-bilayer simulations, two separate profiles (density or dipole) centered around each of the membranes were calculated and then averaged. The reported normalized water dipole orientation corresponds to the cosine of the angle between the water dipole and *z* axis (one of the outputs of gmx h2order analysis) multiplied by the number density of water oxygens, both first calculated as a function of *z*-distance from the center of the membrane. For APL, *D*_P_*_−_*_P_, and the P–N vector tilt, the error was estimated using block averaging.

## Results

### Optimal Parameters for the Flat-Bottom Potential

We first simulated a salt solution with a flat-bottom potential acting on the ions and applied to the plane *z* = 0. The parameters *r* and *k* were systematically varied, and the numbers of ion crossings across the flat-bottom potential plane were extracted for each parameter pair. The results in Table 2 indicate that both *k* and *r* affect the ionic permeability. For the minimum perturbation, combinations of (*r*=0.1 nm, *k*=10,000 kJ*·*mol*^−^*^1^*·*nm*^−^*^2^) and (0.3, 1000) resulted in no crossings, but to play it save, we decided to proceed with the pair of (0.3, 10,000) shown in boldface.

**Table 2:** The number of ion crossings across the flat-bottom potential in the salt solution simulations with different values of ***k*** and ***r***.

| $r$ [nm] | $k$ [kJ·mol<sup>-1</sup>·nm<sup>-2</sup>] | | | |
| --- | --- | --- | --- | --- |
|  | 100 | 1000 | 10,000 | 100,000 |
| 0.1 | 37530 | 19564 | 0 | 0 |
| 0.2 | 27116 | 216 | 0 | 0 |
| 0.3 | 13948 | 0 | **0** | 0 |
| 0.4 | 5054 | 0 | 0 | 0 |
| 0.5 | 1161 | 0 | 0 | 0 |

The inclusion of a flat-bottom potential at *z*_ref_ = 0 pushes ions away from a region 2 *× r* wide and thus increases the ion density elsewhere. Thus, to maintain the properties of the unperturbed bulk away from the flat-bottom restraints, some extra water needs to be included in the system. If the flat-bottom potential was a step function, this region would simply be 2 *× r* wide, and the amount of extra water could be estimated as

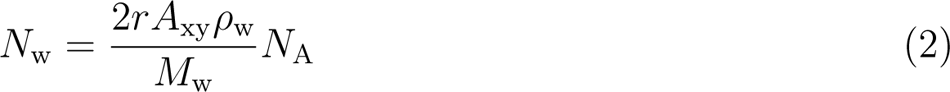

where *N*_w_ is the number of extra water molecules, *A*_xy_ is the area of the flat-bottom potential plane (given that the flat-bottom potential acts along *z* axis), *ρ*_w_ is the water density, *M*_w_ is the molar mass of water (18.016 kg/mol), and *N*_A_ is the Avogadro constant. Yet, the quadratic shape of the flat-bottom potential prevents us from making such assumptions. Thus, we performed further simulations with the salt solution to with different amounts of extra water and with the optimal *k* and *r* values (0.3 nm, 10,000 kJ*·*mol*^−^*^1^*·*nm*^−^*^2^) from the above analysis. We extracted the density profiles of Na^+^ as a function of the *z* coordinate (see Fig. S1 for the profiles), and calculated the average densities in the unperturbed region. These densities are shown in Fig. 2 and compared to the density from an unperturbed system.

**Figure 2:**
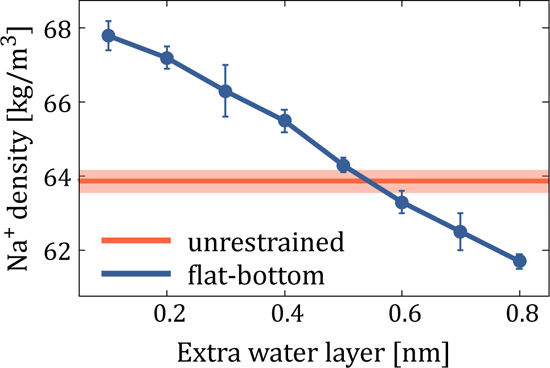
The density of Na^+^ in the unperturbed region as a function of the thickness of the extra water layer is shown in blue. The optimal values of *r*=0.3 nm, *k*=10,000 kJ*·*mol*^−^*^1^*·*nm*^−^*^2^ are used. The error bars show the standard deviation of the density in the bulk region. The red line and the shaded red area show the corresponding average and standard deviation for an unperturbed salt solution, respectively.

The perturbation of the ion distribution in the vicinity of the flat-bottom potential is evident in Fig. S1. However, this is not an issue in typical simulations involving a lipid bilayer since the flat-bottom potential would typically be a few nanometers away from the water–membrane interface and thus screened well by the solvent. The profiles in Fig. 2 suggest that despite the shape of the flat-bottom potential being quadratic, the optimal amount of water to be added is very close to 2 *× r* (here, 0.6 nm). For simplicity, we have used this value in all the bilayer simulations.

Note that, in principle, all our considerations regarding the exact amount of the additional solvent are relevant only to match the ionic concentration in our flat-bottom and double-bilayer setups. Since the entire idea of this work is to promote the usage of the flat-bottom simulations *instead of* double-bilayer approach, the main take-home message from this benchmarking is that correct ionic/solvent concentration should be reported from flat-bottom simulations, taking into account the excluded volume due to ions or other solutes. If the flat-bottom approach is applied to larger molecules (*e.g.*, peptides, drugs, or proteins), the exclusion might need to be correspondingly adjusted. As a rule of thumb, the molecule should be able to sample all orientations within the non-excluded volume and have the desired concentration therein. Often having a specific concentration in the simulation is not crucial—say in the binding of a protein onto the membrane surface—and can be converted for comparison with an experimental value post-simulation. In case concentrations are of more importance, the amount of additional water can be optimized using the cheap and quick solvent simulations following our approach or by using the approximate Eq. (2).

### Lipid Bilayer Simulations with Flat-Bottom Restraints

With the optimal parameters for the flat-bottom potential being established, we applied them to simulations of lipid bilayers with differing solvent environments on their two sides. For comparison, the same bilayers and solvent environments were modeled with the double-bilayer approach. Initially, we checked the simulations where solutions of only monoatomic ions (Na^+^, K^+^, Ca^2+^, Mg^2+^, and Cl*^−^*) were present in the system, *i.e.*, “POPC-NaK-”, “POPC-Xtra-”, and “Plasma-NaK-”, see Table 1. We first characterized the overall dimensions of the bilayer using two common properties, the area per lipid (APL) and membrane thickness defined as the inter-leaflet distance between the phosphorus positions (*D*_P_*_−_*_P_, see Methods). The values of APL and *D*_P_*_−_*_P_ extracted from flat-bottom and double-bilayer setups are shown in Table 3.

**Table 3:** Comparison of the membrane properties (area per lipid (APL), membrane thickness ***D*_P_*_−_*_P_**, and the tilt angle of the P–N vector) calculated from simulations with flat-bottom (FB) and double-bilayer (DB) setups. For asymmetric membranes, the two values of APL are given for the inner and outer leaflets, respectively. The P–N tilt angle is calculated for phosphatidylcholine lipids only and also reported for the two leaflets separately. The error estimate shows the standard error obtained using block averaging.

| System | APL [$\text{\AA}^2$] (inner/outer) | $D_{P-P}$ [nm] | P–N tilt [$^\circ$] (inner/outer) |
| --- | --- | --- | --- |
| POPC-NaK-FB | 64.6±0.1 | 3.89±0.00 | 69.2±0.2 / 68.5±0.0 |
| POPC-NaK-DB | 64.7±0.0 | 3.89±0.00 | 69.2±0.1 / 68.6±0.0 |
| POPC-Xtra-FB | 64.3±0.0 | 3.91±0.00 | 68.5±0.1 / 67.3±0.1 |
| POPC-Xtra-DB | 64.4±0.0 | 3.90±0.00 | 68.7±0.0 / 67.1±0.0 |
| Plasma-NaK-FB | 73.7±0.1 / 75.3±0.1 | 4.63±0.01 | 67.3±0.2 / 66.4±0.3 |
| Plasma-NaK-DB | 73.8±0.1 / 75.4±0.1 | 4.62±0.01 | 67.6±0.2 / 66.5±0.1 |

As Table 3 demonstrates, the bilayer dimensions are indistinguishable between the two used approaches (FB and DB). The addition of divalent cations into the system in the “Xtra” systems leads to a noticeable decrease in APL, and this is reproduced by both approaches. The decrease in APL is coupled to an increase in *D*_P_*_−_*_P_ due to the low compressibility of lipids, and again both approaches provide very similar values. At the level of individual lipids, the tilt angle of the P–N vector depends on the presence of ions and thus shows different values between the two membrane leaflets exposed to different salts. Still, the values in these leaflets are again essentially identical for the flat-bottom and double-bilayer approaches. Similarly, the deuterium order parameters characterizing acyl chain conformations are also indistinguishable between the two approaches, as demonstrated in Fig. S2 in the Supplementary Information (SI). These results highlight that the structure of the bilayer is unaffected by the use of the flat-bottom approach instead of the more computationally demanding double-bilayer one.

Having established that lipids are unaffected by the use of flat-bottom potential, we next looked into the behavior of the solvent. The number density profiles for ions, water, and phosphorus atoms of phospholipids are shown in Fig. 3 with markers and solid lines used for the flat-bottom and double-bilayer approaches, respectively. The excellent agreement between these data indicates that the properties of the solvent are not perturbed by the flat-bottom potential. This is true despite the very different affinities of the ions towards the lipid head groups. Na^+^ shows a density peak at the POPC membrane interface, whereas such a peak is absent for K^+^ (Fig. 3A). Similarly, Ca^2+^ is enriched at the membrane–water interface of the POPC membrane, whereas Mg^2+^ is not (Fig. 3B). In the complex membrane setup, the cytosolic leaflet contains multiple lipid types with charged headgroups, leading to the strong adsorption of K^+^ therein (Fig. 3C). In all studied cases, the density profiles of water and lipid head group phosphorus are also indistinguishable for the flat-bottom and double-bilayer approaches. We acknowledge that the binding of ions heavily depends on the used force field,^50, 51^ but in this work, we contemplate only the agreement between the flat-bottom and double-bilayer approaches.

**Figure 3:**
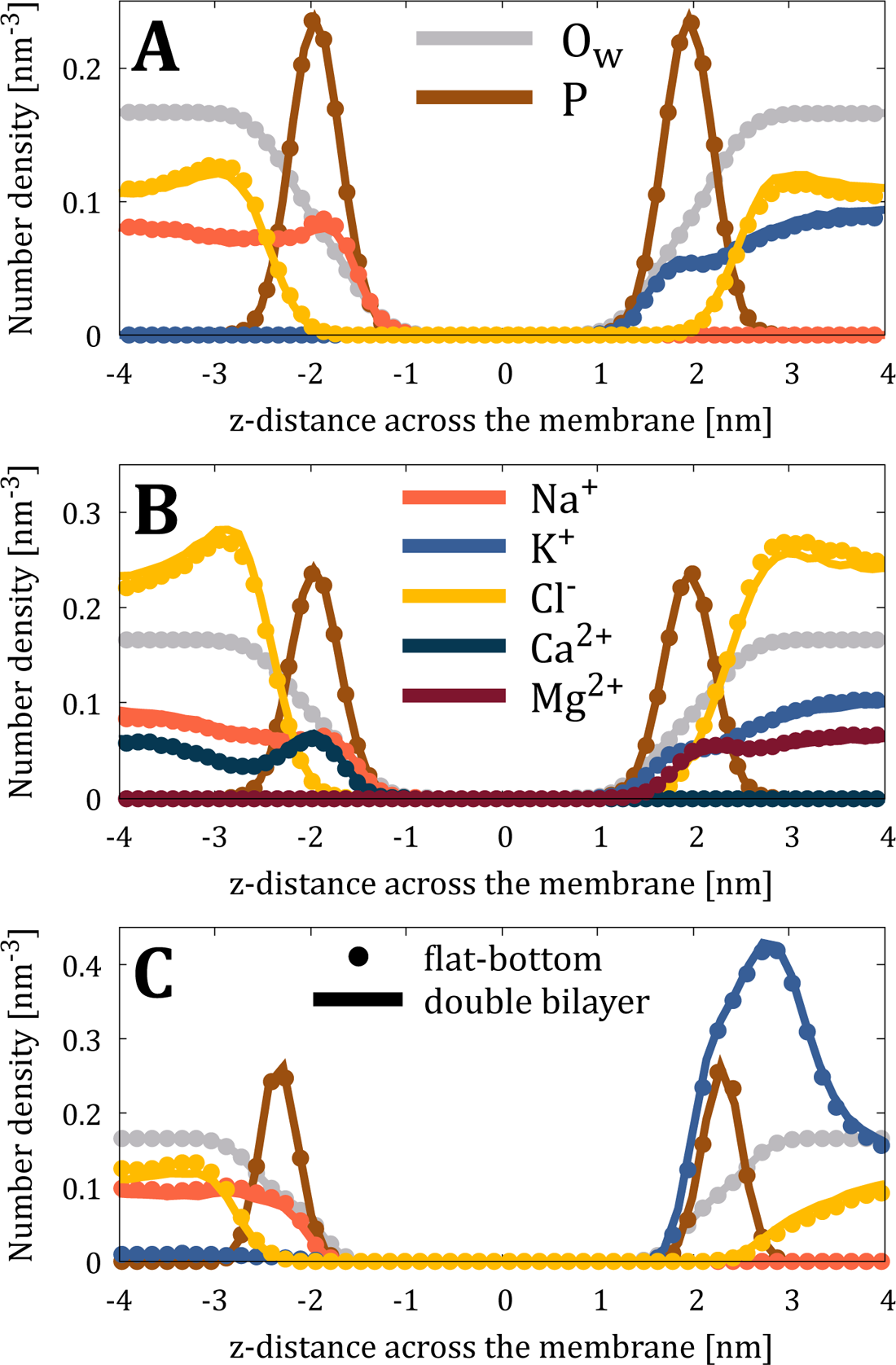
The number density profiles of lipid phosphorus atoms (P, scaled down by a factor of 10 for clarity), water oxygens (O_w_, scaled down by a factor of 200), and ions from flatbottom (markers) and double-bilayer (solid lines) simulations of (A) “POPC-NaK-”, (B) “POPC-Xtra-”, and (C) “Plasma-NaK-systems”.

Although the ionic densities already suggest that using the flat-bottom approach does not induce any undesired electric fields, we also verified this from the orientation of water molecules. The normalized cosine of the water dipole moment (see Methods) is shown for all simulated systems in Fig. 4. A value of zero indicates randomly oriented water, whereas a positive value of the cosine indicates that water molecules are oriented with their hydrogens pointing towards increasing *z* and *vice versa*. The presence of zwitterionic lipid head groups and the adsorbing ions lead to orientational preference close to the membrane interface. However, all such preferences are lost at a distance of *∼*4 nm from the membrane center. Importantly, the profiles are again indistinguishable between the flat-bottom and double-bilayer approaches, strongly suggesting that simulations with the former correctly capture the electrostatic properties of the systems without causing any artifacts, which could arise in ill-designed membrane simulation setups,^52^ *e.g.*, when only cations are restrained by the flat-bottom (see SI).

**Figure 4:**
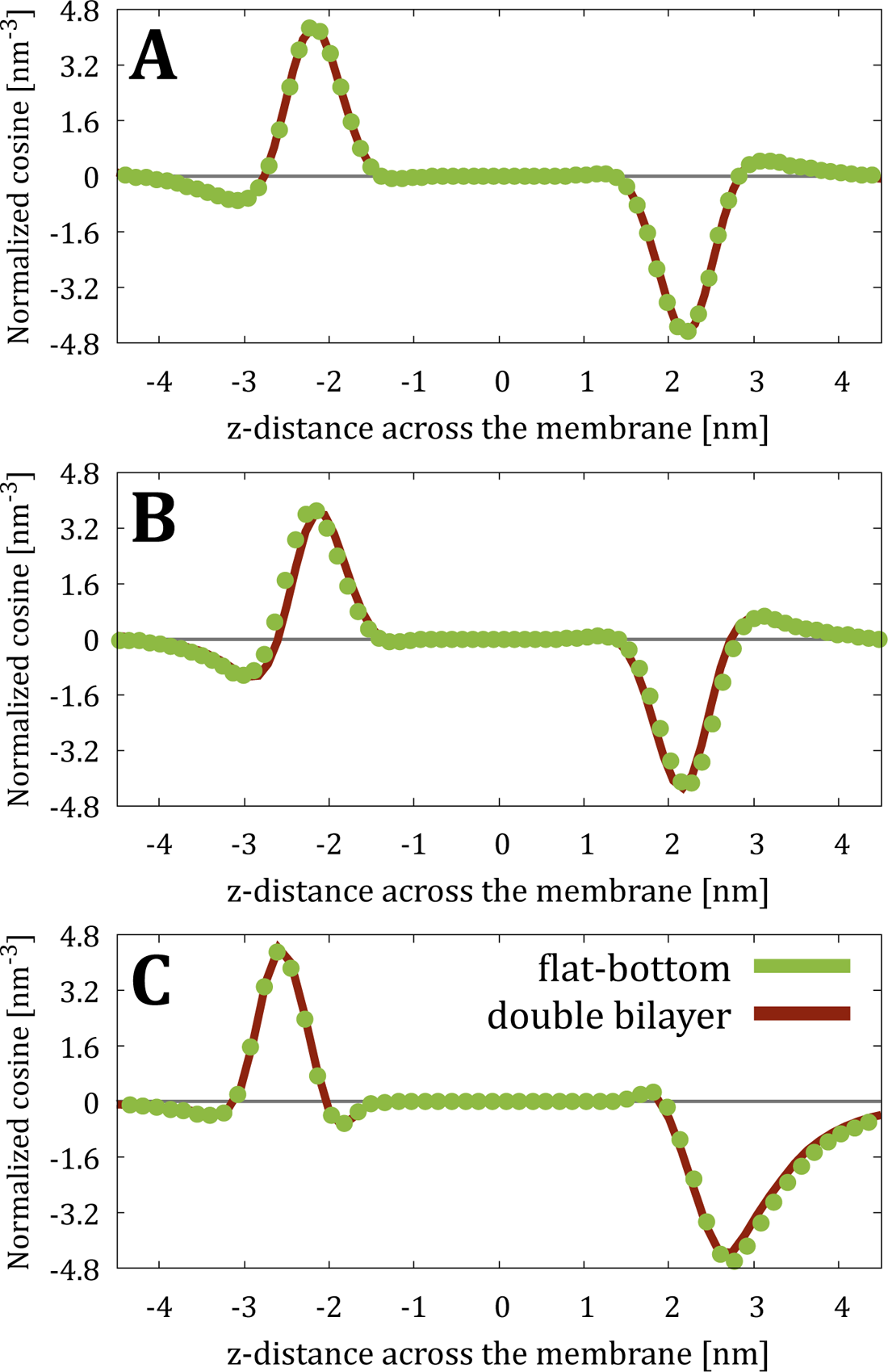
Normalized water orientation from flat-bottom and double-bilayer simulations of (A) “POPC-NaK-”, (B) “POPC-Xtra-”, and (C) “Plasma-NaK-” systems calculated as the cosine of the angle between water dipole and *z* axis multiplied by the number density of water oxygens.

While the results above seem promising, it must be noted that in all cases, there is charge neutrality between the two solvent environments maintained by flat-bottom potentials. While the osmotic coefficients of the solutions on both sides are not necessarily equal, the difference is so small that there is no tendency for the bilayer to drift along its normal (due to the osmotic gradient) to change the volumes and, thus, the concentrations of these solvent environments. This is demonstrated in Fig. S3 in the SI. This figure also shows that when a non-negligible ion imbalance is coupled to the flat-bottom approach, the bilayer rapidly drifts to compensate for this effect, while the flat-bottom potential maintains the ions on the two sides of the membrane. However, this means that the concentrations of the ions might deviate from their initial values. A similar effect would eventually also balance the osmotic pressures of the two water compartments in a double-bilayer setup. However, in this case, the equilibration would be limited by the slow permeation of water through the bilayers, and thus it might not be evident during the typical timescales of atomistic simulations and actually could be revealed sooner in flat-bottom simulations. Finally, modeling extreme cases, *e.g.*, when ions are present on only one side of the membrane, is not recommended in either case, see SI for details. We thereby urge the community to be highly cautious when modeling substantially different solutions at two sides of lipid membranes regardless of the used approach.

### Adsorption of Peptides in Flat-Bottom Simulations

Once we verified the viability of the flat-bottom approach on simpler systems containing only a lipid bilayer and monoatomic ions, we tested its applicability to more complex environments, which involve the adsorption of peptides to lipid membranes. We have chosen nona-arginine (R9) as an example cationic peptide, which can adsorb to both essentially charge-neutral outer leaflet of the “Plasma” membrane (due to specific preferences of R9 peptides to PC lipids^32^) and notably negatively charged inner leaflet (due to attractive electrostatic forces). We tested two situations: (i) when R9 molecules are present only on the extracellular side of the membrane, “Plasma-Asym-”; and (ii) when R9 molecules are present on both sides of the membrane, “Plasma-Pept-”, assuming that some of them entered the cell since R9 are known cell-penetrating peptides.^29, 32^ Note that the design of these simulation setups, Table 1, is purely demonstrative in order to check the agreement between flat-bottom and double-bilayer simulations.

Fig. 5 shows the number density profiles for ions, water, lipid phosphorus atoms, and heavy atoms of R9 peptides. We can observe the excellent agreement between flat-bottom and double-bilayer simulations, including capturing the adsorption of R9 peptides to both leaflets, depletion of Na^+^ and K^+^ cations and even water from the membrane in the presence of R9 (c.f. Fig. 3), and pronounced accumulation of Cl*^−^* anions close to the membrane to compensate for the accumulation of positively charged R9 peptides. All other properties we previously checked for simpler systems (APL, *D*_P_*_−_*_P_, the tilt angle of the P–N vector, deuterium order parameters, and water dipole orientation) are also perfectly reproduced in flat-bottom simulations as compared to double-bilayer setup, see Table S2 and Figs. S4, S5, and S6 in the SI. The perceptible deviations are observed only for the tilt angle, which is expected given the sensitivity of this measure to the adsorption of peptides and only 64 PC lipids being present in the system, *i.e.*, longer simulation times are required for better convergence.

**Figure 5:**
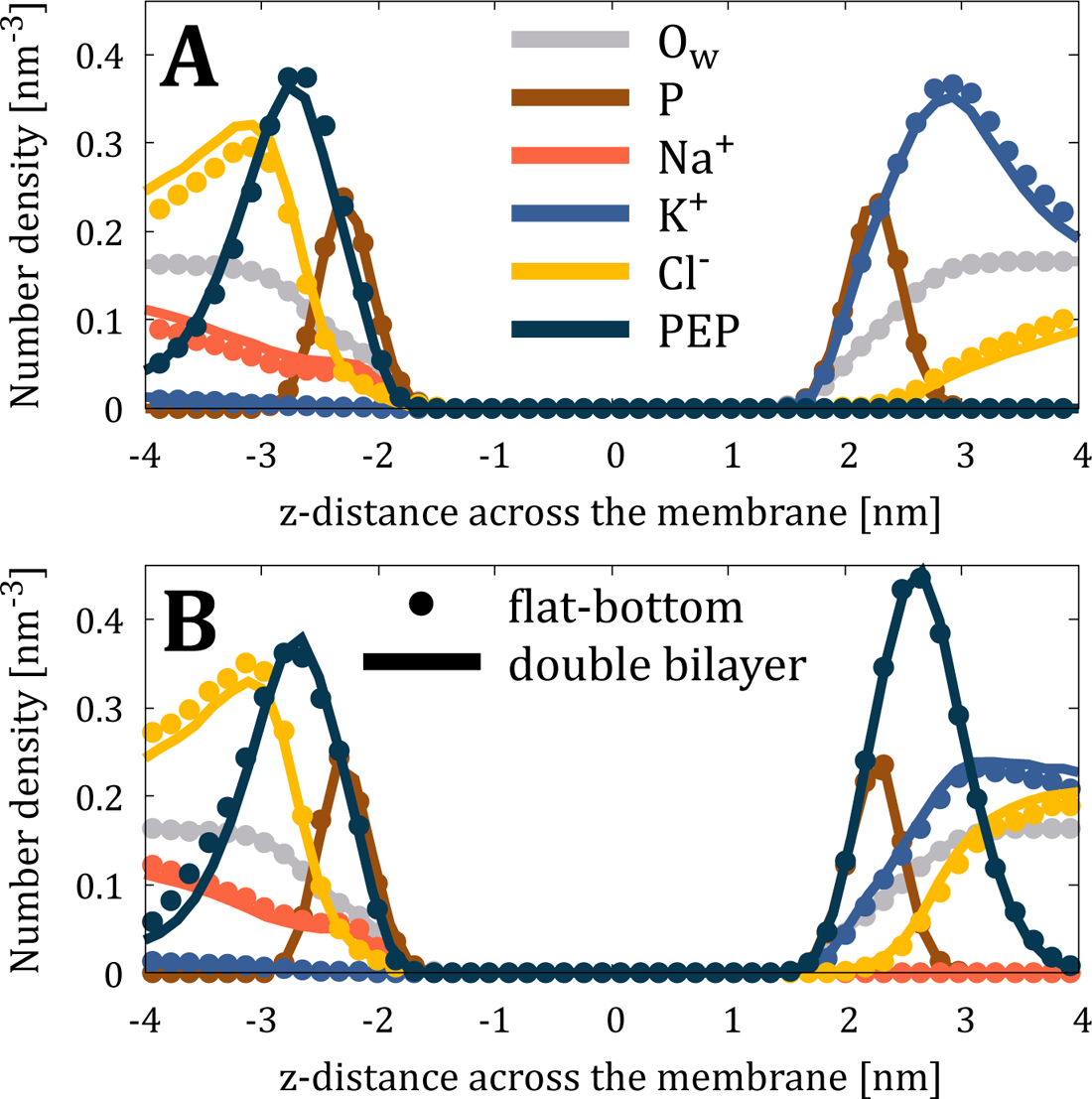
The number density profiles of lipid phosphorus atoms (P, scaled down by a factor of 10 for clarity), water oxygens (O_w_, scaled down by a factor of 200), heavy atoms of nona-arginine peptides (PEP, scaled down by a factor of 10), and ions from flat-bottom (markers) and double-bilayer (solid lines) simulations of (A) “Plasma-Asym-” and (B) “Plasma-Pept-” systems.

### Computational Efficiency of the Flat-Bottom Approach

Finally, it is worth estimating the efficiency of the flat-bottom approach compared to that of the more conventional double-bilayer approach. While the flat-bottom approach generally requires half the particles of the double-bilayer one, it does not lead to a 2-fold speedup in practice. First, a little bit of additional water is required to maintain the correct ionic concentration on the unperturbed solvent region. Secondly, the flat-bottom potential itself needs to be included in the Hamiltonian, thus slightly increasing the computational cost. To estimate the speedup obtained, we benchmarked the flat-bottom and double-bilayer simulation systems on two typical hardware setups: a workstation with a powerful GPU and a CPU-based supercomputer (see SI for more detailed descriptions). As shown in Table 1, the speedup is generally *≈*70% for the workstation with a GPU and 50–80% for a CPU-based supercomputer for the used system sizes. These values were extremely consistent across three repeats. Importantly, for a significantly larger simulation system with 2304 lipids and 576,000 water molecules (dimensions of 27*×*27*×*28 nm^3^), the relative speed of the flat-bottom approach was found to be *≈*2 on both the workstation with a GPU and the CPU-based supercomputer. These results indicate that the flat-bottom approach always leads to a significant reduction in the required computational resources and that it scales particularly well for large simulation systems.

### Summary of the Protocol

Here, we provide step-by-step guidelines on how to set up simulations with the flat-bottom restraints on ions with GROMACS. We also provide example input files in the SI, and all the simulation input and output files used in this work are freely available in the Zenodo repository at DOI:XX

1. **Membrane setup:** If the membrane is set up in CHARMM-GUI,^33^ the extra solvent can be easily provided as an input (“water thickness”), and our analyses suggest that an amount of *r* on each side is sufficient (here 0.3 nm). If you already have the membrane set up, the box can be extended in *z*, the membrane re-centered with gmx editconf, and the extra solvent added with gmx solvate. With other tools such as Packmol^53^ or insane (in the case of coarse-grained simulations),^54^ the box size can be provided as an input.
2. **Ion addition:** As the flat-bottom potential prevents ions from crossing the *z*_ref_ = 0 plane, they must be initially correctly positioned. We have simply concatenated structure (.gro) files containing the ions at desired locations with the membrane structure, followed by a thorough energy minimization. Alternatively, the ions can be inserted using gmx insert-molecules, whose -ip option can be used to limit each ion type to a specific region in the simulation box. With Packmol,^53^ the ions can readily be placed in desired regions either when the system is being built or as an additional step.
3. **Restraint file:** The reference coordinates *z*_ref_ are read from an additional .gro file provided to grompp with the -r handle. Thus, the *z* coordinates of all ions must be set to 0 in this file. This can be readily achieved with a simple script modifying the fields of a regular .gro file of the system or using the replace functionality of a text editor on the desired column. For example, using awk, this is achieved with the command “awk ‘NR==1 print NR==2 natoms=$0; print NR>2 && NR<natoms print substr($0,1,38) “ 0.000” NR==natoms+3 print’ IN > OUT”, where IN and OUT are the structure file for the system and the restraint file with all *z* coordinates equal to zero, respectively. Note that the .gro file format is sensitive to formatting, which is preserved by this command.
4. **Topology setup:** Finally, the flat-bottom restraints need to be added to the molecule-types of all the ions (or other molecules) to which the restraints are applied. The [position restraints] statement used in this work for an atom number N of the moleculetype reads “N 2 5 −0.3 10000”, where we apply a flat-bottom restraint (2) of type 5 (layer, *z*) with an *r* value of 0.3 nm (minus sign inverses the potential as desired) and with a *k* value of 10,000 kJ*·*mol*^−^*^1^*·*nm*^−^*^2^. In the case of atomic ions, each moleculetype consists of only one atom, and thus N = 1. In the case of peptides, we applied the restraints only to heavy atoms, *i.e.*, non-hydrogens, to minimize the external bias while still preventing the crossing of periodic boundary conditions. Another way to introduce position restraints is using PLUMED,^55^ a software patch compatible with GROMACS and primarily designed to perform metadynamics calculations and feed forces to the MD engine. PLUMED also allows applying a restraint on the center of mass of a molecule instead of each selected atom separately. This could be a more suitable solution for complex molecules like peptides and proteins. However, a decrease in speed performance is anticipated in this case, so we recommend using PLUMED only when its other features are used simultaneously, and thus the flat-bottom restraints do not become the speed-limiting factor.
5. **Generating run input files:** gmx grompp will generate a warning unless the refcoord scaling is set to “all” or “com”. These options define how the *z*_ref_ values are scaled based on box size fluctuations. However, since the used *z*_ref_ values are 0 in our approach, they are not scaled regardless of which option is chosen, if there are no other applied restraints in the system. If this is not the case (*e.g.*, when lipid atoms are restrained during the equilibration), the reference positions will be mutually scaled when “com” option is chosen. Yet, the “all” option functions as expected without any effect on the positioning of the flat-bottom restraints at *z*_ref_ = 0.

## Conclusions

In this paper, we have described an efficient protocol for simulating a lipid bilayer interacting with two distinct solvent environments enabled by using flat-bottom restraints. This approach can be used to facilitate faster sampling or to model physiologically relevant ion gradients across lipid membranes. Using flat-bottom restraints instead of a more conventional double-bilayer setup provides at least a 1.5-fold speedup with small lipid bilayer simulations, and the speedup increases to 2-fold with larger system sizes.

Here, we have carefully evaluated the parameters for the flat-bottom potential that lead to the least possible perturbation to the simulated systems. With this optimal parameter set, we demonstrated that the structures of both the lipid bilayer and solvents are indistinguishable between the flat-bottom and double-bilayer approaches. In our examples, we considered various lipid and solvent compositions, including ones with charged peptides, yet the approach can readily be applied to other molecules of physiological or biotechnological interest, such as drugs and other small molecules. The flat-bottom method has no force-field dependencies and can be easily applied in both all-atom and coarse-grained simulations. While the flat-bottom approach is suitable for studies of structural properties under equilibrium conditions—such as the distribution of ions and the adsorption geometry of proteins at the membrane–water interface—care must be taken when applying it to non-equilibrium simulations, such as those tackling permeation events. Additionally, both the flat-bottom and double-bilayer approaches should be applied only in cases where the osmotic pressures of the two solvent environments do not differ significantly, yet the issues become apparent significantly quicker with the former.

## Acknowledgement

We thank CSC–IT Center for Science for computing resources. DB acknowledges support from the project “National Institute of Virology and Bacteriology (Program EXCELES, ID Project No. LX22NPO5103) – Funded by the European Union – Next Generation EU”. MJ thanks the Academy of Finland (Postdoctoral researcher grant no. 338160) and the Emil Aaltonen foundation for funding. We thank Balazs Fabian and Timothée Rivel for testing and optimizing the preparation and simulation protocols, including simplifying our example awk command.

## Supporting Information Available

Methodology for the estimation of speedup in the flat-bottom approach. Density profiles of Na^+^ in aqueous salt solution simulations with different amounts of extra water in flat-bottom simulations. Methodology and results for systems with unrestrained Cl*^−^* ions. Methodology and results for systems with a large charge imbalance across the leaflets. Composition of the membrane in the “Plasma” simulations. Comparison of the deuterium order parameters of the lipid acyl chains between the flat-bottom and double-bilayer setups. Drift of the membrane center of mass in the flat-bottom simulations. Additional results from simulations with R9 peptides.

## References

1. Rothman, J. E.; Lenard, J. Membrane asymmetry. Science 1977, 195, 743–753.

2. Van Meer, G.; Voelker, D. R.; Feigenson, G. W. Membrane lipids: where they are and how they behave. *Nature Rev*. Mol. Cell Biol. 2008, 9, 112–124.

3. Lorent, J.; Levental, K.; Ganesan, L.; Rivera-Longsworth, G.; Sezgin, E.; Doktorova, M.; Lyman, E.; Levental, I. Plasma membranes are asymmetric in lipid unsaturation, packing and protein shape. Nat. Chem. Biol. 2020, 16, 644–652.

4. Fadeel, B.; Xue, D. The ins and outs of phospholipid asymmetry in the plasma membrane: Roles in health and disease. Crit. Rev. Biochem. Mol. Biol 2009, 44, 264–277.

5. Daleke, D. L. Regulation of transbilayer plasma membrane phospholipid asymmetry. J. Lipid Res. 2003, 44, 233–242.

6. Lodish, H.; Berk, A.; Zipursky, S. L.; Matsudaira, P.; Baltimore, D.; Darnell, J. Molecular Cell Biology. 4th edition; WH Freeman, 2000.

7. Enkavi, G.; Javanainen, M.; Kulig, W.; Ŕog, T.; Vattulainen, I. Multiscale simulations of biological membranes: The challenge to understand biological phenomena in a living substance. Chem. Rev. 2019, 119, 5607–5774.

8. Marrink, S. J.; Corradi, V.; Souza, P. C.; Inǵolfsson, H. I.; Tieleman, D. P.; Sansom, M. S. Computational modeling of realistic cell membranes. Chem. Rev. 2019, 119, 6184–6226.

9. Blumer, M.; Harris, S.; Li, M.; Martinez, L.; Untereiner, M.; Saeta, P. N.; Carpenter, T. S.; Inǵolfsson, H. I.; Bennett, W. Simulations of asymmetric membranes illustrate cooperative leaflet coupling and lipid adaptability. Front. Cell Dev. Biol. 2020, 8, 575.

10. Inǵolfsson, H. I.; Melo, M. N.; Van Eerden, F. J.; Arnarez, C.; Lopez, C. A.; Wassenaar, T. A.; Periole, X.; De Vries, A. H.; Tieleman, D. P.; Marrink, S. J. Lipid organization of the plasma membrane. J. Am. Chem. Soc. 2014, 136, 14554–14559.

11. Inǵolfsson, H. I.; Carpenter, T. S.; Bhatia, H.; Bremer, P.-T.; Marrink, S. J.; Lightstone, F. C. Computational lipidomics of the neuronal plasma membrane. Biophys. J. 2017, 113, 2271–2280.

12. Park, S.; Beaven, A. H.; Klauda, J. B.; Im, W. How tolerant are membrane simulations with mismatch in area per lipid between leaflets? J. Chem. Theory Comput. 2015, 11, 3466–3477.

13. Doktorova, M.; Weinstein, H. Accurate in silico modeling of asymmetric bilayers based on biophysical principles. Biophys. J. 2018, 115, 1638–1643.

14. Miettinen, M. S.; Lipowsky, R. Bilayer membranes with frequent flip-flops have tensionless leaflets. Nano Lett. 2019, 19, 5011–5016.

15. Varma, M.; Deserno, M. Distribution of cholesterol in asymmetric membranes driven by composition and differential stress. Biophys. J. 2022, 121, 4001–4018.

16. Sachs, J. N.; Crozier, P. S.; Woolf, T. B. Atomistic simulations of biologically realistic transmembrane potential gradients. J. Chem. Phys. 2004, 121, 10847–10851.

17. Gurtovenko, A. A. Asymmetry of lipid bilayers induced by monovalent salt: Atomistic molecular-dynamics study. J. Chem. Phys. 2005, 122, 244902.

18. Lee, S.-J.; Song, Y.; Baker, N. A. Molecular dynamics simulations of asymmetric NaCl and KCl solutions separated by phosphatidylcholine bilayers: potential drops and structural changes induced by strong Na^+^-lipid interactions and finite size effects. Biophys. J. 2008, 94, 3565–3576.

19. Denning, E. J.; Woolf, T. B. Double bilayers and transmembrane gradients: A molecular dynamics study of a highly charged peptide. Biophys. J. 2008, 95, 3161–3173.

20. Darden, T.; York, D.; Pedersen, L. Particle mesh Ewald: An N*·*log(N) method for Ewald sums in large systems. J. Chem. Phys. 1993, 98, 10089–10092.

21. Wennberg, C. L.; Murtola, T.; Hess, B.; Lindahl, E. Lennard-Jones lattice summation in bilayer simulations has critical effects on surface tension and lipid properties. J. Chem. Theory Comput. 2013, 9, 3527–3537.

22. Melcr, J.; Bonhenry, D.; Timr, S.; Jungwirth, P. Transmembrane potential modeling: Comparison between methods of constant electric field and ion imbalance. J. Chem. Theory Comput. 2016, 12, 2418–2425.

23. Kasparyan, G.; Hub, J. S. Equivalence of charge imbalance and external electric fields during free energy calculations of membrane electroporation. J. Chem. Theory Comput. 2023, 19, 2676–2683.

24. Bilkova, E.; Pleskot, R.; Rissanen, S.; Sun, S.; Czogalla, A.; Cwiklik, L.; Ŕog, T.; Vattulainen, I.; Cremer, P. S.; Jungwirth, P., et al. Calcium directly regulates phosphatidylinositol 4,5-bisphosphate headgroup conformation and recognition. J. Am. Chem. Soc. 2017, 139, 4019–4024.

25. Duboue-Dijon, E.; Javanainen, M.; Delcroix, P.; Jungwirth, P.; Martinez-Seara, H. A practical guide to biologically relevant molecular simulations with charge scaling for electronic polarization. J. Chem. Phys. 2020, 153, 050901.

26. Luo, Y.; Roux, B. Simulation of osmotic pressure in concentrated aqueous salt solutions. J. Phys. Chem. Lett. 2010, 1, 183–189.

27. Sinelnikova, A.; Spoel, D. v. d. NMR refinement and peptide folding using the GRO-MACS software. J. Biomol. NMR 2021, 75, 143–149.

28. Bischoff, M.; Biriukov, D.; P̌redota, M.; Roke, S.; Marchioro, A. Surface Potential and Interfacial Water Order at the Amorphous TiO_2_ Nanoparticle/Aqueous Interface. J. Phys. Chem. C 2020, 124, 10961–10974.

29. Allolio, C.; Magarkar, A.; Jurkiewicz, P.; Baxová, K.; Javanainen, M.; Mason, P. E.; Šachl, R.; Cebecauer, M.; Hof, M.; Horinek, D., et al. Arginine-rich cell-penetrating peptides induce membrane multilamellarity and subsequently enter via formation of a fusion pore. Proc. Natl. Acad. Sci. 2018, 115, 11923–11928.

30. Abraham, M. J.; Murtola, T.; Schulz, R.; Páll, S.; Smith, J. C.; Hess, B.; Lindahl, E. GROMACS: High performance molecular simulations through multi-level parallelism from laptops to supercomputers. SoftwareX 2015, 1, 19–25.

31. Páll, S.; Zhmurov, A.; Bauer, P.; Abraham, M.; Lundborg, M.; Gray, A.; Hess, B.; Lindahl, E. Heterogeneous parallelization and acceleration of molecular dynamics simulations in GROMACS. J. Chem. Phys. 2020, 153, 134110.

32. Nguyen, M. T. H.; Biriukov, D.; Tempra, C.; Baxova, K.; Martinez-Seara, H.; Evci, H.; Singh, V.; Šachl, R.; Hof, M.; Jungwirth, P., et al. Ionic strength and solution composition dictate the adsorption of cell-penetrating peptides onto phosphatidylcholine membranes. Langmuir 2022, 38, 11284–11295.

33. Jo, S.; Lim, J. B.; Klauda, J. B.; Im, W. CHARMM-GUI Membrane Builder for mixed bilayers and its application to yeast membranes. Biophys. J. 2009, 97, 50–58.

34. Wu, E. L.; Cheng, X.; Jo, S.; Rui, H.; Song, K. C.; Dávila-Contreras, E. M.; Qi, Y.; Lee, J.; Monje-Galvan, V.; Venable, R. M., et al. CHARMM-GUI membrane builder toward realistic biological membrane simulations. J. Comput. Chem. 2014, 35, 1997– 2004.

35. Lee, J.; Cheng, X.; Swails, J. M.; Yeom, M. S.; Eastman, P. K.; Lemkul, J. A.; Wei, S.; Buckner, J.; Jeong, J. C.; Qi, Y. et al. CHARMM-GUI input generator for NAMD, GROMACS, AMBER, OpenMM, and CHARMM/OpenMM simulations using the CHARMM36 additive force field. J. Chem. Theory Comput. 2016, 12, 405–413.

36. Klauda, J. B.; Venable, R. M.; Freites, J. A.; O’Connor, J. W.; Tobias, D. J.; Mondragon-Ramirez, C.; Vorobyov, I.; MacKerell Jr, A. D.; Pastor, R. W. Update of the CHARMM all-atom additive force field for lipids: validation on six lipid types. J. Phys. Chem. B 2010, 114, 7830–7843.

37. Lim, J. B.; Rogaski, B.; Klauda, J. B. Update of the cholesterol force field parameters in CHARMM. J. Phys. Chem. B 2012, 116, 203–210.

38. Jorgensen, W. L.; Chandrasekhar, J.; Madura, J. D.; Impey, R. W.; Klein, M. L. Comparison of simple potential functions for simulating liquid water. J. Chem. Phys. 1983, 79, 926–935.

39. Durell, S. R.; Brooks, B. R.; Ben-Naim, A. Solvent-induced forces between two hydrophilic groups. J. Phys. Chem. 1994, 98, 2198–2202.

40. Huang, J.; Rauscher, S.; Nawrocki, G.; Ran, T.; Feig, M.; De Groot, B. L.; Grubmüller, H.; MacKerell, A. D. CHARMM36m: an improved force field for folded and intrinsically disordered proteins. Nat. Methods 2017, 14, 71–73.

41. Nencini, R.; Tempra, C.; Biriukov, D.; Poĺak, J.; Ondo, D.; Heyda, J.; Ollila, S. O.; Javanainen, M.; Martinez-Seara, H. Prosecco: polarization reintroduced by optimal scaling of electronic continuum correction origin in MD simulations. Biophys. J. 2022, 121, 157a.

42. Páll, S.; Hess, B. A flexible algorithm for calculating pair interactions on SIMD architectures. Comput. Phys. Commun. 2013, 184, 2641–2650.

43. Essmann, U.; Perera, L.; Berkowitz, M. L.; Darden, T.; Lee, H.; Pedersen, L. G. A smooth particle mesh Ewald method. J. Chem. Phys. 1995, 103, 8577–8593.

44. Nośe, S. A unified formulation of the constant temperature molecular dynamics methods. J. Chem. Phys. 1984, 81, 511–519.

45. Hoover, W. G. Canonical dynamics: Equilibrium phase-space distributions. Phys. Rev. A 1985, 31, 1695.

46. Parrinello, M.; Rahman, A. Polymorphic transitions in single crystals: A new molecular dynamics method. J. Appl. Phys. 1981, 52, 7182–7190.

47. Hess, B. P-LINCS: A parallel linear constraint solver for molecular simulation. J. Chem. Theory Comput. 2008, 4, 116–122.

48. Hess, B.; Bekker, H.; Berendsen, H. J.; Fraaije, J. G. LINCS: a linear constraint solver for molecular simulations. J. Comput. Chem. 1997, 18, 1463–1472.

49. Miyamoto, S.; Kollman, P. A. SETTLE: An analytical version of the SHAKE and RATTLE algorithm for rigid water models. J. Comput. Chem. 1992, 13, 952–962.

50. Catte, A.; Girych, M.; Javanainen, M.; Loison, C.; Melcr, J.; Miettinen, M. S.; Monticelli, L.; Määattä, J.; Oganesyan, V. S.; Ollila, O. S., et al. Molecular electrometer and binding of cations to phospholipid bilayers. Phys. Chem. Chem. Phys. 2016, 18, 32560–32569.

51. Antila, H.; Buslaev, P.; Favela-Rosales, F.; Ferreira, T. M.; Gushchin, I.; Javanainen, M.; Kav, B.; Madsen, J. J.; Melcr, J.; Miettinen, M. S. et al. Headgroup structure and cation binding in phosphatidylserine lipid bilayers. J. Phys. Chem. B 2019, 123, 9066–9079.

52. Kopec, W.; Gapsys, V. Periodic boundaries in Molecular Dynamics simulations: why do we need salt? bioRxiv 2022, 2022.10.18.512672.

53. Martínez, L.; Andrade, R.; Birgin, E. G.; Martínez, J. M. PACKMOL: A package for building initial configurations for molecular dynamics simulations. J. Comput. Chem. 2009, 30, 2157–2164.

54. Wassenaar, T. A.; Inǵolfsson, H. I.; Bockmann, R. A.; Tieleman, D. P.; Marrink, S. J. Computational lipidomics with *insane*: A versatile tool for generating custom membranes for molecular simulations. J. Chem. Theory Comput. 2015, 11, 2144–2155.

55. Tribello, G. A.; Bonomi, M.; Branduardi, D.; Camilloni, C.; Bussi, G. PLUMED 2: New feathers for an old bird. Comput. Phys. Commun. 2014, 185, 604–613.

