## Supporting Information for "Efficient Simulations of Membrane and Solvent Asymmetry With Flat-Bottom Restraints"

### Supplementary Methods

#### Estimation of Relative Speed

The speed gain obtained using the flat-bottom approach compared to the double-bilayer one was estimated using both a workstation computer and a supercomputer. The workstation was equipped with an AMD Ryzen 9 5950X 16-core processor, an nVidia GeForce RTX3080 Ti GPU, and 128 GB of RAM. On the supercomputer, four nodes were used, and each was equipped with two AMD Rome 7H12 64-core processors and 256 GB memory. No GPUs were used in the calculation on the supercomputer. For the benchmark, we ran all the systems listed in Table 1 in three replicates and used their mean as a representative speed estimate. The first 10,000 steps were discarded due to the PME optimization, after which the speed was recorded for 500,000 steps (1 ns). Finally, we calculated the ratios of the simulation speeds obtained with the flat-bottom and double-bilayer approaches, which are reported in Table 1. For the larger test system containing 2,304 DMPC lipids, 576,000 waters (for the flat-bottom approach, an additional water layer of 0.6 nm was included), and 150 mM NaCl, we used 10 nodes on the supercomputer for our tests.

### Supplementary Results

We provide additional data that compare the results from flat-bottom and double-bilayer simulations and confirm the viability of the former approach. We also additionally discuss how the flat-bottom simulations behave if chloride anions are free of restraints or ions are placed only on one side of the lipid membrane.

#### Chloride Ions Free of Restraints

Since chloride ions do not really interact with lipid membranes, one tempting approach is to include them on both sides of the plasma membrane unrestrained, therefore allowing them to move freely between two solvent environments. We tested this idea on some of our systems and identified several issues with this approach. First, the ionic concentrations of cations and anions become imbalanced since their accessible volume is different, being larger for chloride ions, see density profiles in Fig. S7. This leads to a lower effective concentration of chloride compared to cations. Second, unrestrained chlorides cause artificial water orientation in the vicinity of the flat-bottom excluded volume, Fig. S8. This artifact is the consequence of a little accumulation of cations in this region, which is not balanced by the same accumulation of chloride ions that do not feel the flat-bottom potential. Therefore, we do not recommend using this hybrid approach in flat-bottom restraints, although all membrane and interfacial properties are unaffected by the free motion of chloride ions, see Figs. S7 and S9, and Table S3.

#### Solutes and Ions Only on One Side of the Membrane

Using the flat-bottom approach, one may model the extreme conditions when ions or other solutes are present only on one side of the membrane. In Fig. S10, we show that such system design leads to a large osmotic gradient that pushes the membrane from the middle of the box until it essentially reaches the position of the flat-bottom restrain, *i.e.* the solvent becomes fully available to all ions and solutes present in the system. Interestingly, the same should happen in double-bilayer simulations (potentially leading to the collapse of two bilayers) but does not occur on the timescale of typical membrane simulations due to the slow exchange of water between two solvent compartments.

Table S1: Lipid compositions of the inner and outer leaflets of the realistic membrane[S1] used in simulations of "Plasma-NaK", "'Plasma-Asym", and "Plasma-Pept" setups. The composition is slightly different from the original one due to the availability of corresponding lipids in CHARMM-GUI[S2–S4]

| Lipid | $№_{inner}$ | $№_{outer}$ |
|---|---|---|
| PAPS | 20 | 0 |
| POPC | 6 | 0 |
| POPE | 4 | 0 |
| PDOPE | 12 | 0 |
| SAPE | 6 | 0 |
| DPPC | 4 | 0 |
| SAPI24 | 4 | 0 |
| SAPS | 2 | 2 |
| PLA20 | 18 | 4 |
| PLPC | 14 | 22 |
| PSM | 2 | 18 |
| NSM | 0 | 14 |
| LSM | 0 | 12 |
| SOPC | 0 | 10 |
| PAPC | 0 | 8 |
| CHOL | 57 | 65 |

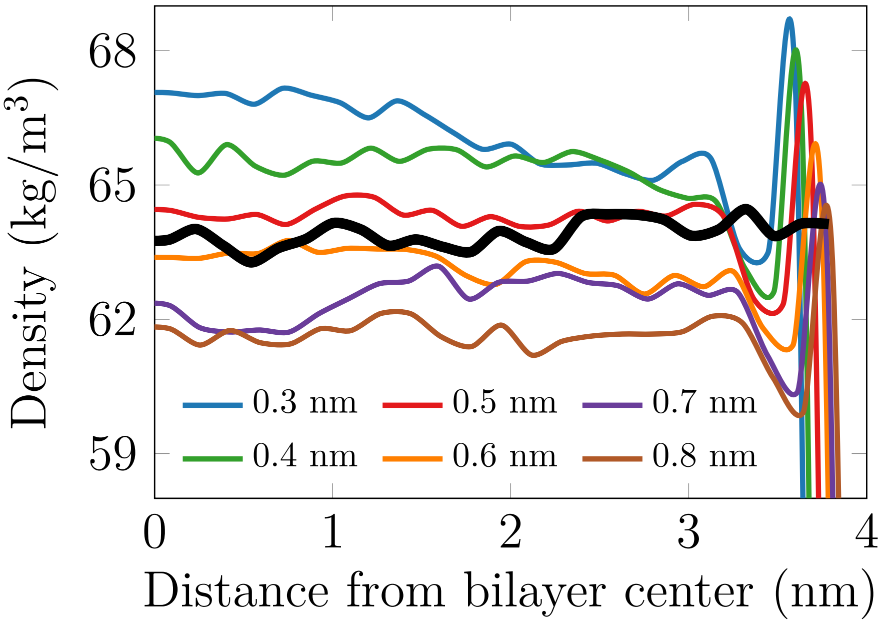


Figure S1: Density profiles for $Na^+$ with the optimal parameters for ($r$=0.3 nm, $k$ =10,000 kJ·mol$^{-1}$·nm$^{-2}$) and with various amounts of extra water added. The solid black line refers to a simulation without flat-bottom restraints.

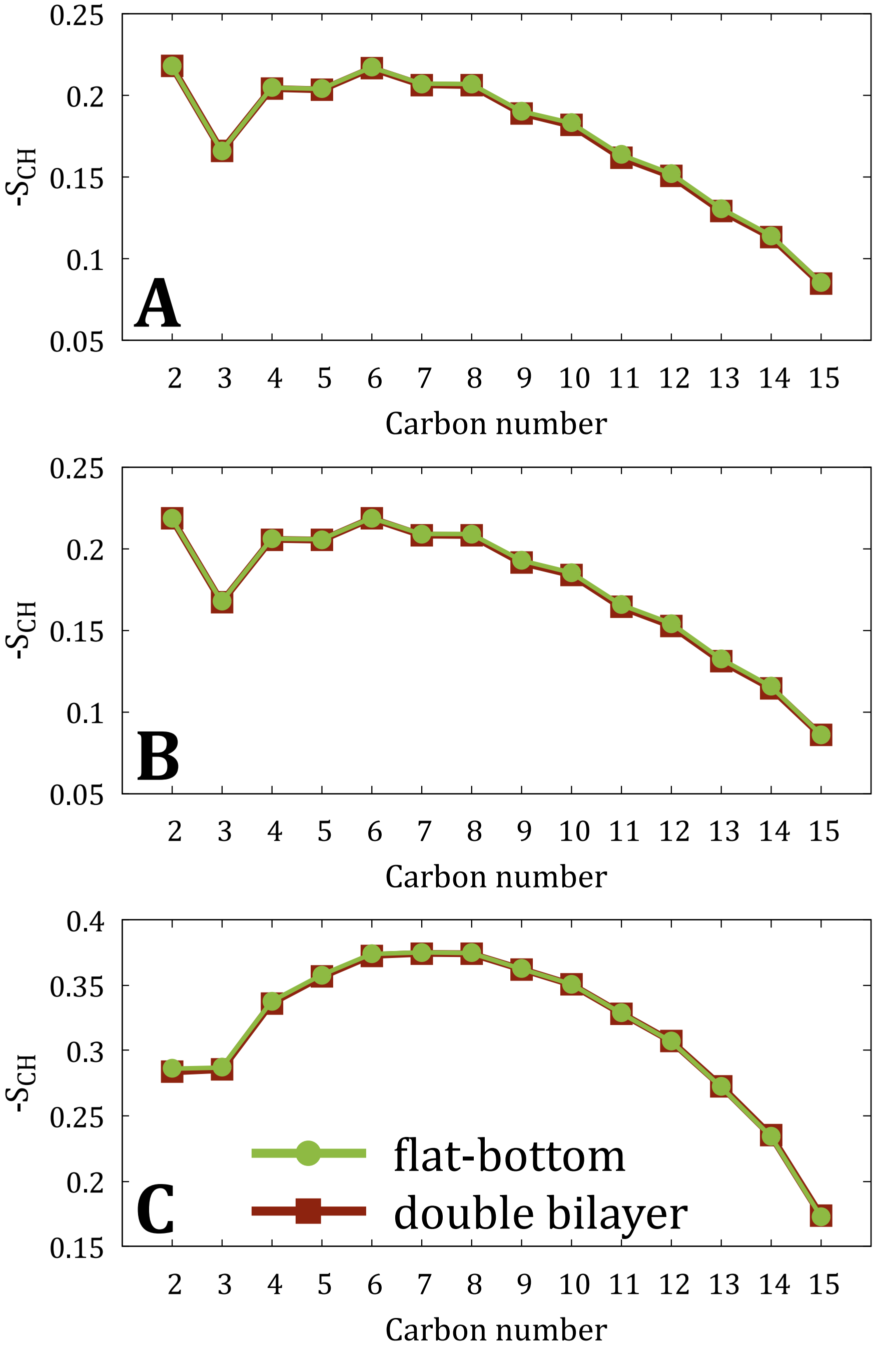


Figure S2: $S_{CH}$ calculated for the 16:0 *sn*-1 acyl chains, *i.e.*, palmitate, present in PAPS, POPC, POPE, PDOPE, DPPC, PLPC, and PAPC lipids from flat-bottom and double-bilayer simulations of (A) "POPC-NaK-", (B) "POPC-Xtra-", and (C) "Plasma-NaK-" systems.

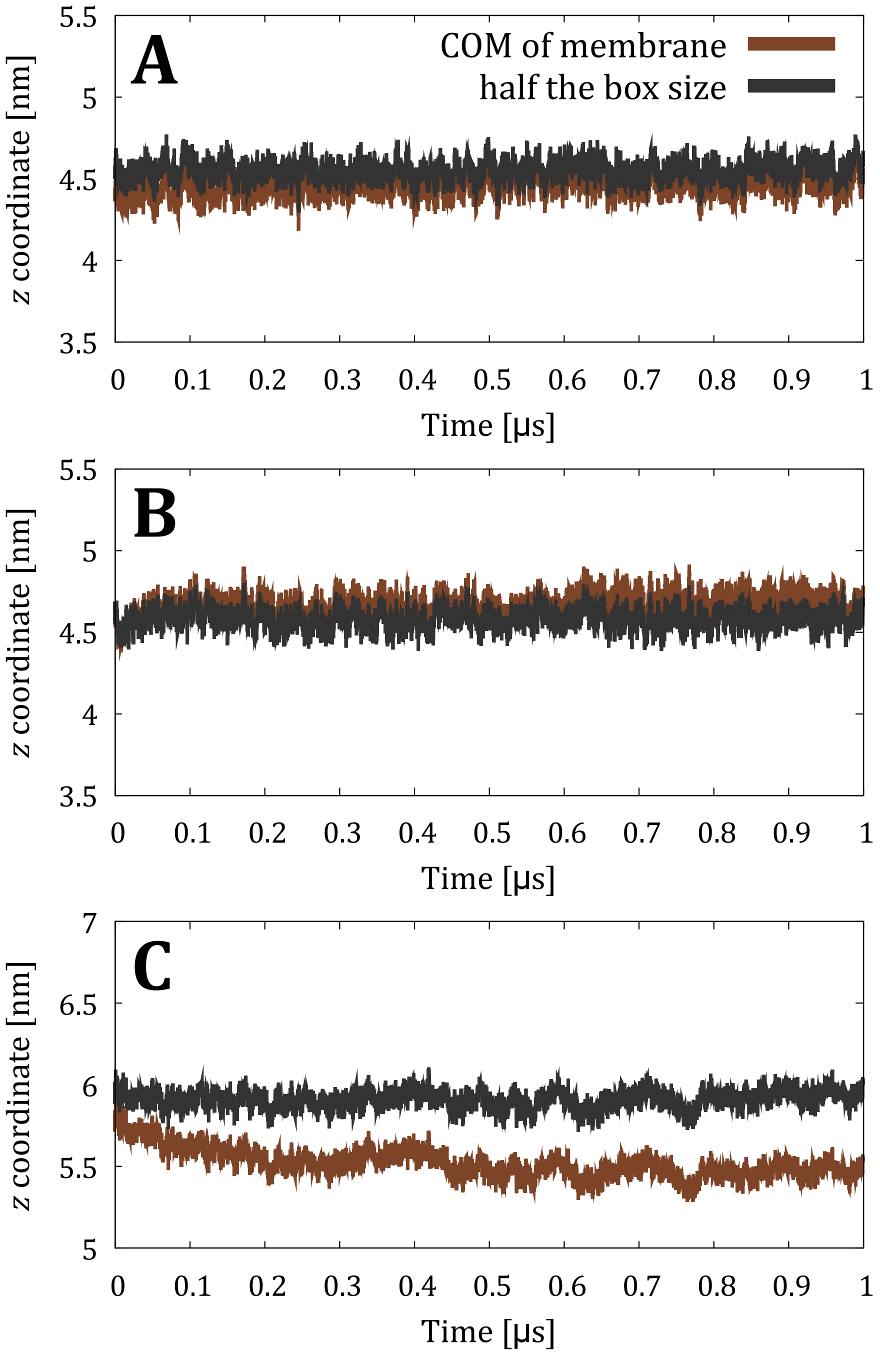


Figure S3: The $z$ coordinate of the center of mass of the membrane compared to the $z$ dimension of the box size as a function of simulation time from flat-bottom simulations of (A) "POPC-NaK-", (B) "POPC-Xtra-", and (C) "Plasma-NaK-" systems.

Table S2: Comparison of the membrane properties (area per lipid (APL), membrane thickness $D_{\mathrm{P-P}}$, and the tilt angle of the P–N vector) calculated from R9 peptides-containing simulations with flat-bottom (FB) and double-bilayer (DB). The two values of APL are given for the inner and outer leaflets, respectively. The P–N tilt angle is calculated for phosphatidylcholine lipids only and also reported for the two leaflets separately. The error estimate shows the standard error obtained using block averaging.

| System | APL [Å$^2$] (inner/outer) | $D_{\mathrm{P-P}}$ [nm] | P–N tilt [°] (inner/outer) |
|---|---|---|---|
| Plasma-Asym-FB | 73.6±0.1 / 75.3±0.1 | 4.53±0.01 | 62.4±0.0 / 59.2±0.4 |
| Plasma-Asym-DB | 73.7±0.1 / 75.3±0.1 | 4.52±0.00 | 62.3±0.2 / 58.2±0.4 |
| Plasma-Pept-FB | 73.0±0.1 / 74.7±0.1 | 4.56±0.01 | 60.4±0.3 / 58.6±0.3 |
| Plasma-Pept-DB | 73.4±0.1 / 75.1±0.1 | 4.54±0.00 | 61.4±0.2 / 63.0±0.3 |

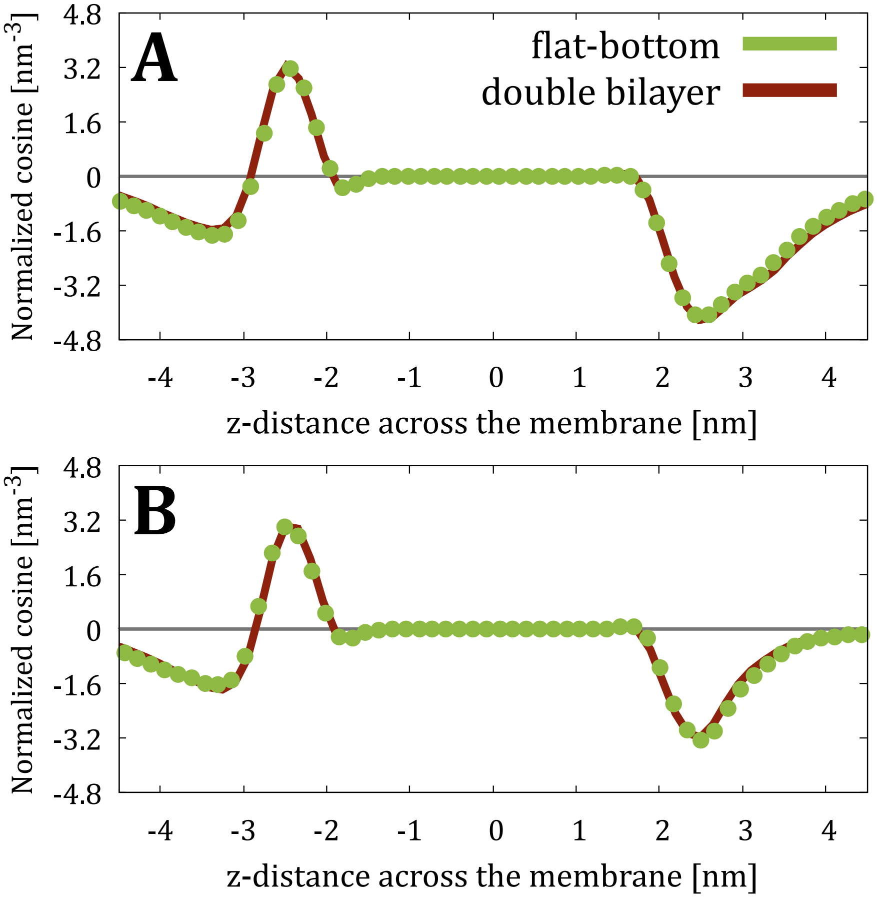


Figure S4: Normalized water orientation from flat-bottom (markers) and double-bilayer (solid lines) simulations of (A) “Plasma-Asym-” and (B) “Plasma-Pept-” systems calculated as the cosine of the angle between water dipole and $z$ axis multiplied by the number density of water oxygens.

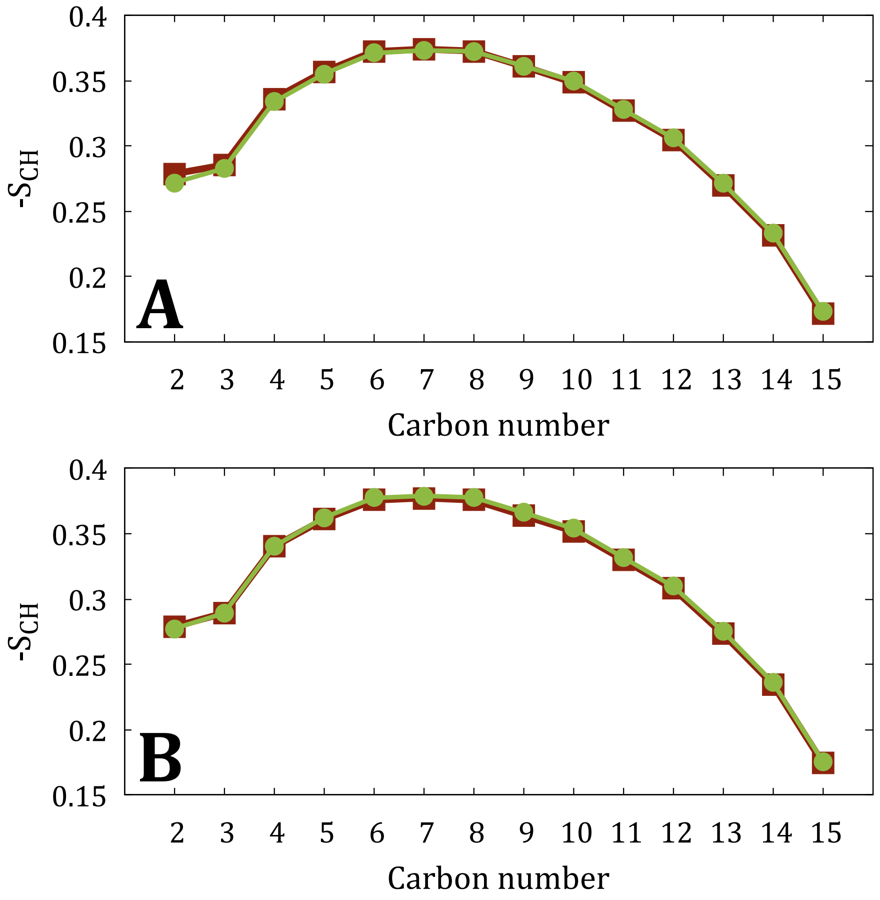


Figure S5: $S_{CH}$ calculated for the 16:0 *sn*-1 acyl chains, *i.e.*, palmitate, present in PAPS, POPC, POPE, PDOPE, DPPC, PLPC, and PAPC lipids from flat-bottom and double-bilayer simulations of(A) "Plasma-Asym-" and (B) "Plasma-Pept-" systems.

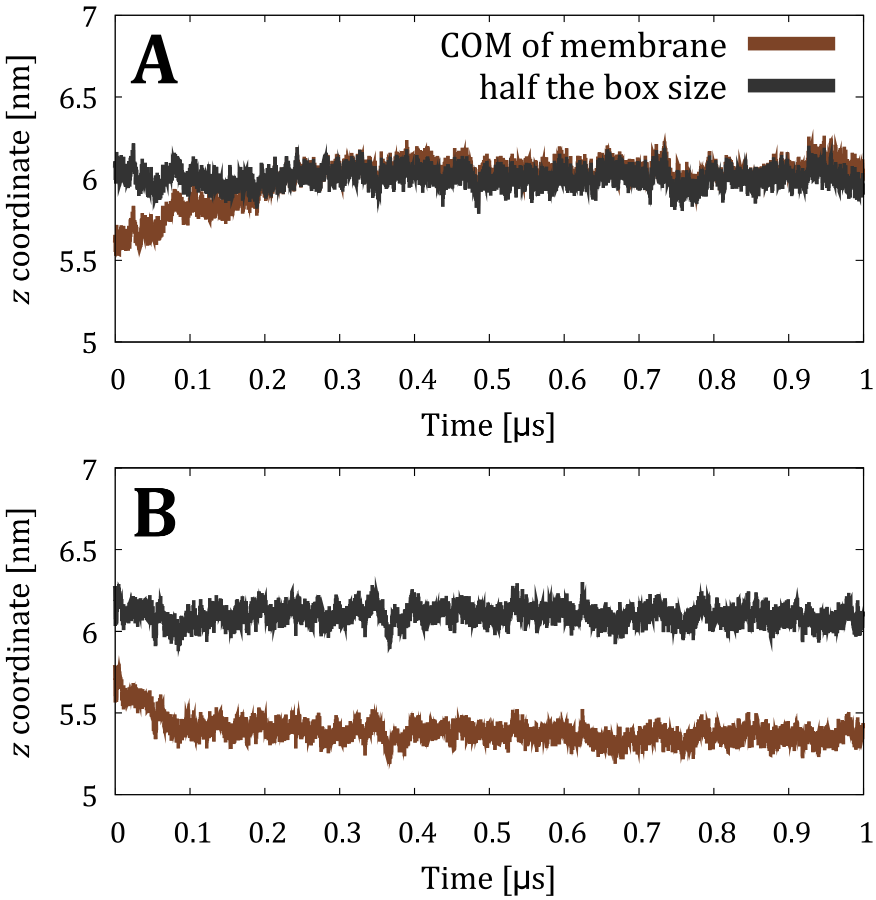


Figure S6: The $z$ coordinate of the center of mass of the membrane compared to the $z$ dimension of the half box size as a function of simulation time from flat-bottom and double-bilayer simulations of(A) "Plasma-Asym-" and (B) "Plasma-Pept-" systems.

Table S3: Comparison of the membrane properties (area per lipid (APL), membrane thickness $D_{P-P}$, and the tilt angle of the P–N vector) calculated from simulations with flat-bottom with free chloride ions (FB-freeCl) and double-bilayer (DB) setups. For asymmetric membranes, the two values of APL are given for the inner and outer leaflets, respectively. The P–N tilt angle is calculated for phosphatidylcholine lipids only and also reported for the two leaflets separately. The error estimate shows the standard error obtained using block averaging.

| System | APL [Å$^2$] (inner/outer) | $D_{P-P}$ [nm] | P–N tilt [°] (inner/outer) |
|---|---|---|---|
| POPC-NaK-FB-freeCl | 64.8±0.0 | 3.88±0.00 | 69.3±0.1 / 68.5±0.1 |
| POPC-NaK-DB | 64.7±0.0 | 3.89±0.00 | 69.2±0.1 / 68.6±0.0 |
| POPC-Xtra-FB-freeCl | 64.4±0.0 | 3.90±0.00 | 68.6±0.1 / 67.4±0.1 |
| POPC-Xtra-DB | 64.4±0.0 | 3.90±0.00 | 68.7±0.0 / 67.1±0.0 |
| Plasma-NaK-FB-freeCl | 73.8±0.1 / 75.5±0.1 | 4.62±0.00 | 67.2±0.2 / 66.5±0.2 |
| Plasma-NaK-DB | 73.8±0.1 / 75.4±0.1 | 4.62±0.01 | 67.6±0.2 / 66.5±0.1 |

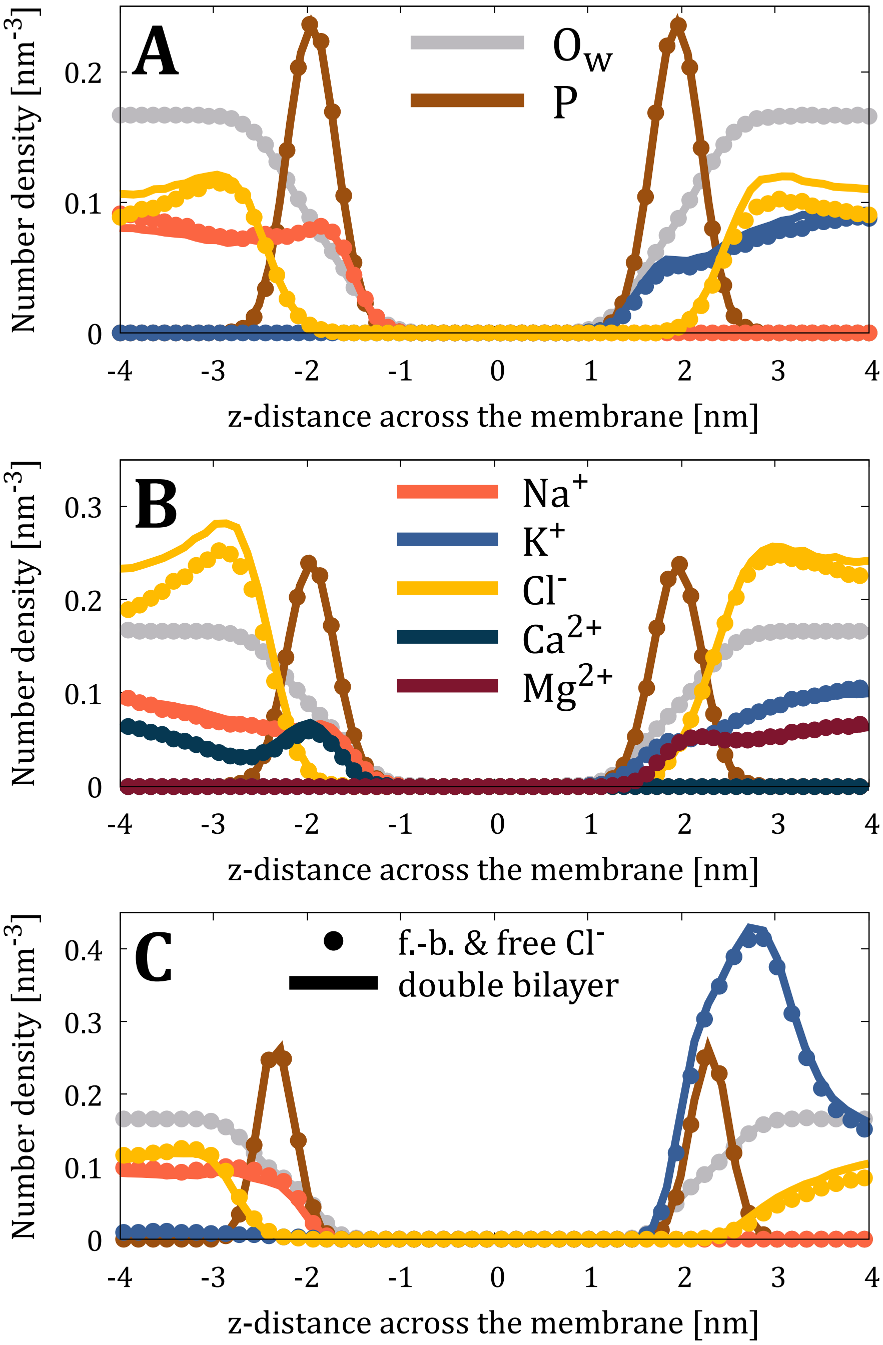


Figure S7: The number density profiles of lipid phosphorus atoms (P, scaled down by a factor of 10 for clarity), water oxygens ($O_w$, scaled down by a factor of 200), and ions from flat-bottom with free chloride ions (markers; "f.-b. & free $Cl^-$") and double-bilayer (solid lines) simulations of (A) "POPC-NaK-", (B) "POPC-Xtra-", and (C) "Plasma-NaK-" systems.

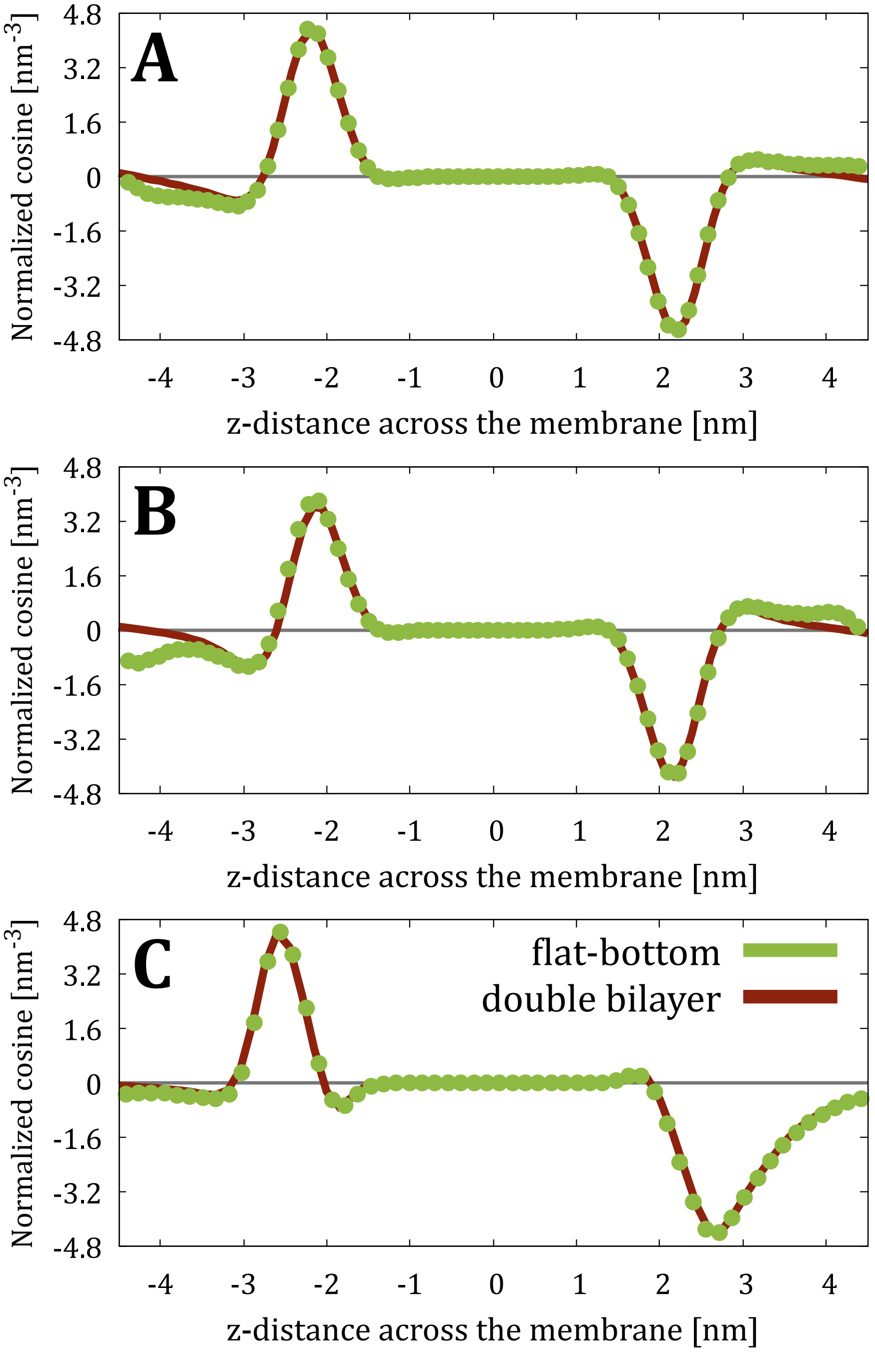


Figure S8: Normalized water orientation from flat-bottom with free chloride ions ("f.-b. & free Cl$^-$") and double-bilayer simulations of (A) "POPC-NaK-", (B) "POPC-Xtra-", and (C) "Plasma-NaK-" systems calculated as the cosine of the angle between water dipole and $z$-axis multiplied by the number density of water oxygens.

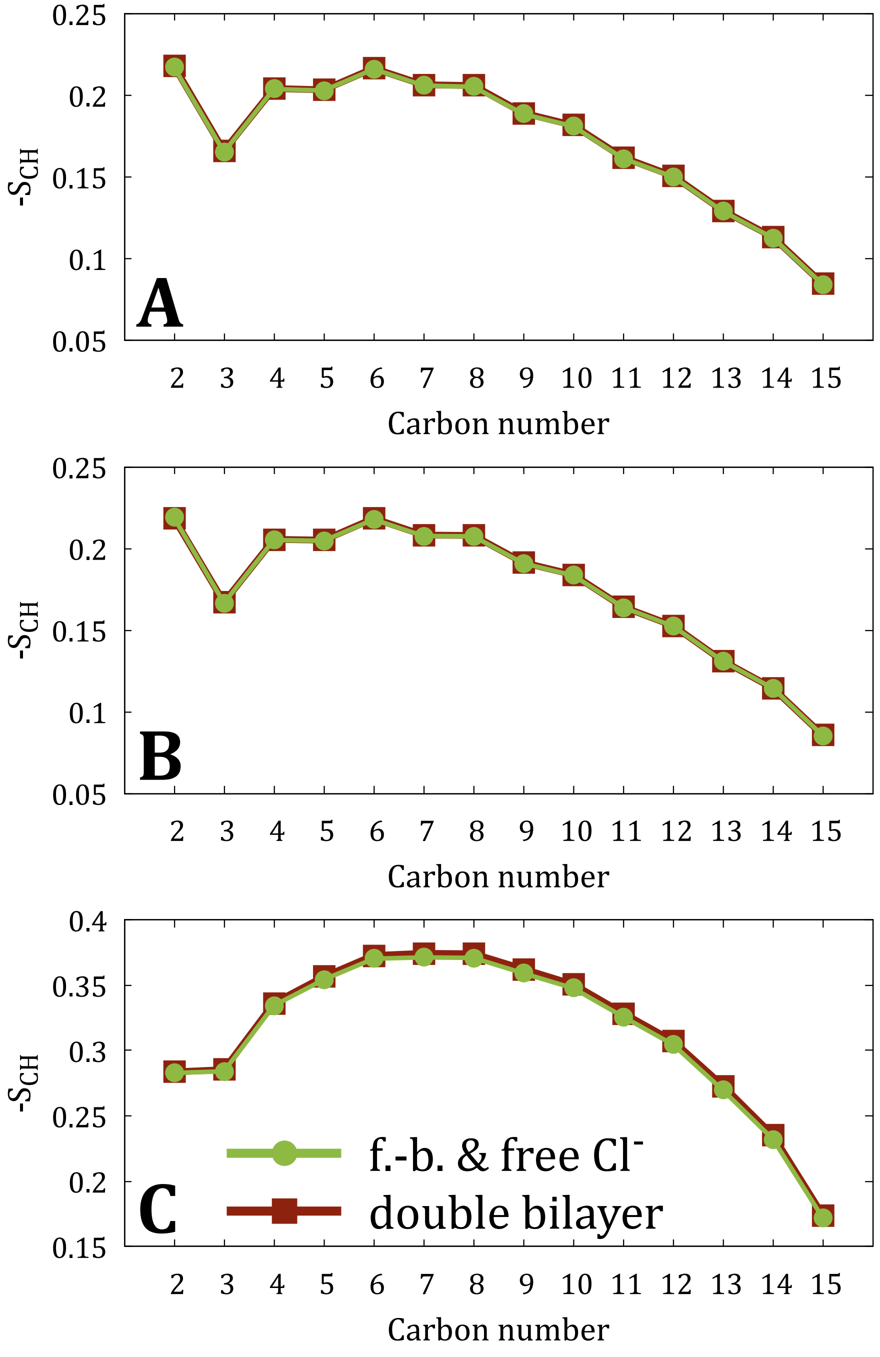


Figure S9: $S_{CH}$ calculated for the 16:0 *sn*-1 acyl chains, *i.e.*, palmitate, present in PAPS, POPC, POPE, PDOPE, DPPC, PLPC, and PAPC lipids from flat-bottom simulations with free chloride ions ("f.-b. & free $Cl^-$") and double-bilayer simulations of (A) "POPC-NaK-", (B) "POPC-Xtra-", and (C) "Plasma-NaK-" systems.

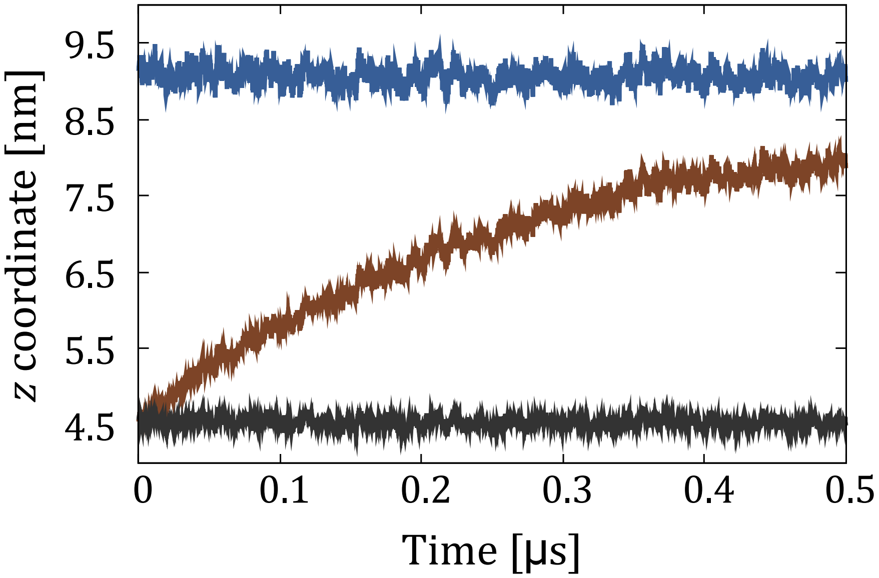


Figure S10: The $z$ coordinate of the center of mass of the membrane compared to the $z$ dimension of the half box size as a function of simulation time from flat-bottom simulations with ions present only on one side of the membrane.

### References


(S1) Lorent, J.; Levental, K.; Ganesan, L.; Rivera-Longsworth, G.; Sezgin, E.; Doktorova, M.; Lyman, E.; Levental, I. Plasma membranes are asymmetric in lipid unsaturation, packing and protein shape. *Nat. Chem. Biol.* **2020**, *16*, 644–652.

(S2) Lee, J.; Cheng, X.; Swails, J. M.; Yeom, M. S.; Eastman, P. K.; Lemkul, J. A.; Wei, S.; Buckner, J.; Jeong, J. C.; Qi, Y.; Jo, S.; Pande, V. S.; Case, D. A.; Brooks, C. L.; MacKerell, A. D. J.; Klauda, J. B.; Im, W. CHARMM-GUI input generator for NAMD, GROMACS, AMBER, OpenMM, and CHARMM/OpenMM simulations using the CHARMM36 additive force field. *J. Chem. Theory Comput.* **2016**, *12*, 405–413.

(S3) Wu, E. L.; Cheng, X.; Jo, S.; Rui, H.; Song, K. C.; Dávila-Contreras, E. M.; Qi, Y.; Lee, J.; Monje-Galvan, V.; Venable, R. M.; Klauda, J. B.; Im, W. CHARMM-GUI membrane builder toward realistic biological membrane simulations. *J. Comput. Chem.* **2014**, *35*, 1997–2004.

(S4) Jo, S.; Lim, J. B.; Klauda, J. B.; Im, W. CHARMM-GUI Membrane Builder for mixed bilayers and its application to yeast membranes. *Biophys. J.* **2009**, *97*, 50–58.
